# Tuning DO:DM ratios modulates MHC class II immunopeptidomes

**DOI:** 10.1101/2021.10.05.463141

**Authors:** Niclas Olsson, Wei Jiang, Lital N. Adler, Elizabeth D. Mellins, Joshua E. Elias

## Abstract

Major histocompatibility complex class II (MHC-II) antigen presentation underlies a wide range of immune responses in health and disease. However, how MHC-II antigen presentation is regulated by the peptide-loading catalyst HLA-DM (DM), its associated modulator, HLA-DO (DO), is incompletely understood. This is due largely to technical limitations: model antigen presenting cell (APC) systems that express these MHC-II peptidome regulators at physiologically variable levels have not been described. Likewise, computational prediction tools that account for DO and DM activities are not presently available. To address these gaps, we created a panel of single MHC-II allele, HLA-DR4-expressing APC lines that cover a wide range of DO:DM ratio states. Using a combined immunopeptidomic and proteomic discovery strategy, we measured the effects DO:DM ratios have on peptide presentation by surveying over 10,000 unique DR4-presented peptides. The resulting data provide insight into peptide characteristics that influence their presentation with increasing DO:DM ratios. These include DM-sensitivity, peptide abundance, binding affinity and motif, peptide length and register positioning on the source protein. These findings have implications for designing improved HLA-II prediction algorithms and research strategies for dissecting the variety of functions that different APCs serve in the body.

**IN BRIEF:** Peptides presented by MHC-II are critical to adaptive immune function. The non-canonical MHC molecules HLA-DM and HLA-DO cooperatively regulate MHC-II function, but how varied DO-to-DM ratios across different APCs and cellular contexts might influence their immunopeptide repertoires is unclear. We address this by measuring cell lines expressing these two proteins spanning a range of relative abundances. We found that peptides could be categorized according to how robustly they were presented at different DO:DM ratios. Importantly, this presentation was only partially linked to predicted affinity to the MHC-II molecule.

**HIGHLIGHTS:**

- Describe MHC-class II peptide repertoires from a unique HLA-DR4 cell line panel with increasing DO:DM ratios.
- Demonstrate striking and divergent changes in MHC-II immunopeptidomes that result from the tuning function of DO:DM.
- These findings bridge gap in understanding and predicting MHC-II antigen presentation.

**GRAPHICAL ABSTRACT:** 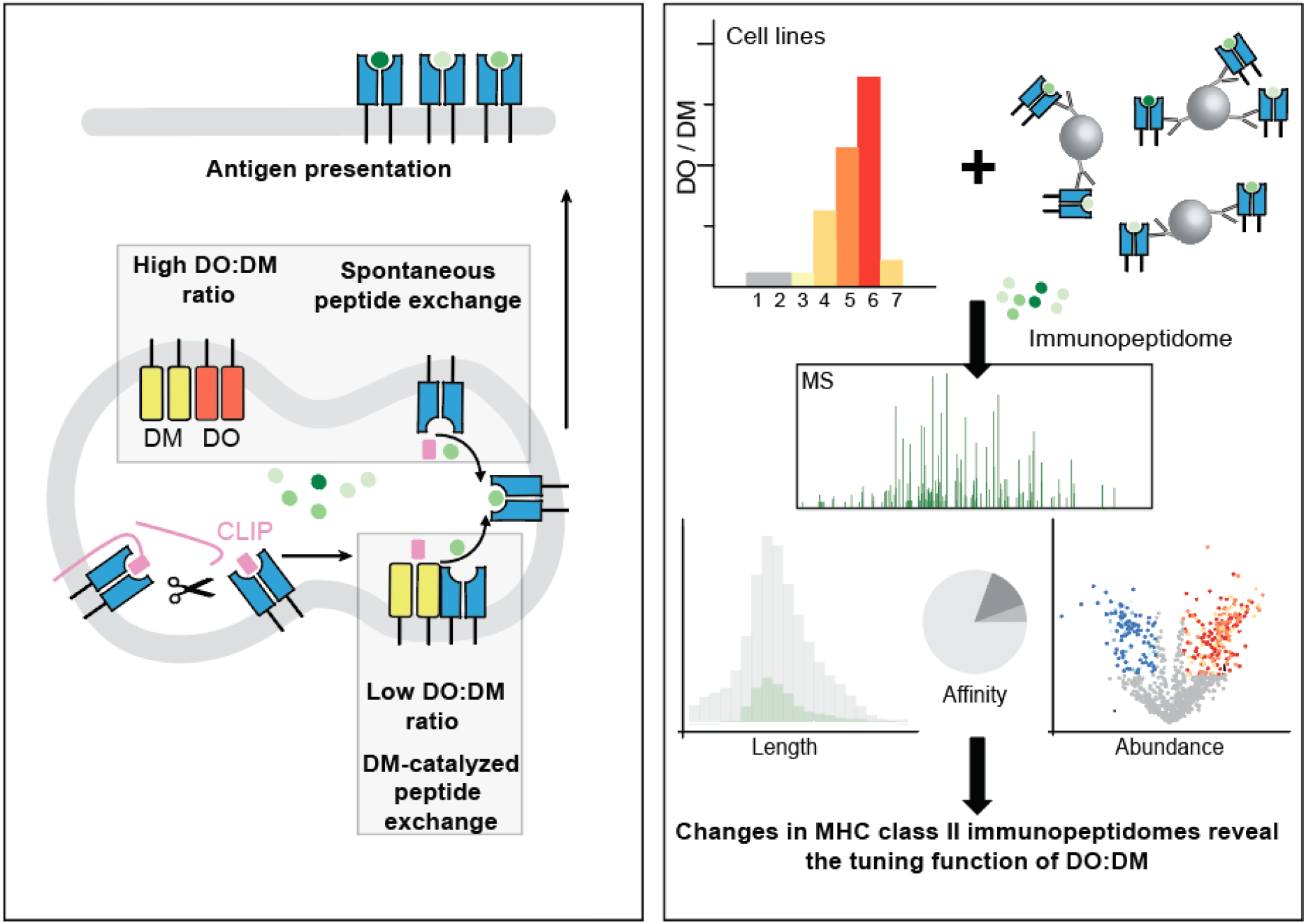

## INTRODUCTION

The immune system develops effector T cell repertoires that are both tolerant of self-proteins and reactive to foreign antigens through selection steps that remove T cell clones bearing high-affinity, self-reactive antigen receptors (1, 2). Professional antigen-presenting cells (APCs) present peptides in complex with major histocompatibility complex class II (pMHC-II), in a process that is essential to CD4+ T cell development and clonal selection. During self-tolerance acquisition in the thymus, medullary thymic epithelia and other APCs present self-pMHC-II where they are encountered by developing CD4+ T, driving their maturation and selection (1). During antigen exposure in the periphery, diverse pMHC-II presented by APCs, like dendritic cells and B cells in the germinal centers (GC) of lymph nodes, expand antigen-reactive mature T cell clones (2). In both circumstances, APCs express a non-classical MHC-II molecule, DO (HLA-DO in human or H2-O in mouse) (3–7). DO is expressed by medullary thymic epithelia, B cells and certain dendritic cell subsets, but not by macrophages (8–10). This restricted expression pattern implies a unique yet critical role for DO in regulating pMHC-II presentation and subsequent CD4+ T-dependent immune responses.

Consistent with DO-regulated immune responses, ectopic overexpression of H2-O in H2-O^neg^ dendritic cells diminished the presentation of certain pMHC-II (11) and prevented diabetogenic T cell activation and subsequent type 1 diabetes symptoms in NOD mice (12). Conversely, H2-O knockout mice spontaneously developed increased T-dependent autoantibody titers but showed delayed humoral immunity when immunized with a model antigen (13). These mice also exhibited increased susceptibility to model autoimmune diseases (14). In contrast, other researchers found that H2-O-deficiency promoted virus-neutralizing antibody production (15). These experiments support the idea that the immunologic consequences of DO’s effects on pMHC-II repertoires are highly context dependent such that DO knockout animals can exhibit autoreactive or anti-microbial responses compared to DO-sufficient animals. Physiologically, DO expression is downregulated in naïve or memory B cells entering GC to recruit T cell help (5, 6, 16). DO downregulation also occurs in certain dendritic cells following their exposure to maturation stimuli (8, 9). Thus, changes in DO levels influence a wide range of adaptive immune responses.

Recently, we proposed a mechanistic model of how DO regulates MHC-II peptide presentation, considering both molecular and cellular assessments of DO function (17). DO functions through its pH-dependent association with DM (18–21), a peptide exchange catalyst that selects for stable pMHC-II (22, 23), although not all high affinity peptides are resistant to DM. The DM selection process is termed editing. We also found that the presented peptidome can be highly sensitive to DO regulation, as small changes in DO:DM stoichiometry (the DO-to-DM ratio is denoted as DO:DM) can create substantial shifts in pMHC-II levels (17). As noted above, several studies have shown that DO knockout or ectopic expression in murine or human cells influences certain pMHC-II or the entire MHC-II peptidome (11, 14, 24–30). However, no study has yet measured how pMHC-II repertoires change between graded DO:DM differences, such as those that exist among APC types, or those that progressively occur upon APC activation (8, 17).

Here, we performed a detailed mass-spectrometric (MS) analysis on the peptidomes associated with an MHC-II human leucocyte antigen allele, HLA-DR4 expressed by model B cell lines, by jointly considering the related proteomic information. These cell lines cover a wider range of DO:DM stoichiometry than the previously used DO+ versus DO-states. We demonstrated striking and divergent changes in HLA-II peptidomes that result from the tuning function of DO:DM. We also described a knock-down example that illustrates how DO downregulation would affect such tuning effect on the peptidome. Using these MS analyses, we identified several key characteristics associated with DO:DM-tuned pMHC-II presentation. Our findings thus will advance research in DMDO-regulated antigen presentation, improve computational prediction models of MHC-II antigen presentation, and suggest strategies in management of T-dependent immune disorders.

## EXPERIMENTAL PROCEDURES

### Cell line construction

The previously described cell lines included in this work are T2 (MHC-II^null^DO^null^DM^null^ TxB hybrid), T2DR4 (DO^null^DM^null^), T2DR4DM and two clonal cell lines, T2DR4DMDO++ (1C3) and T2DR4DMDO+++ (2D7) (17, 21). Cell lines with graded levels of DO:DM were constructed as follows. A T2DR4DMDO parent line was constructed by transfecting the pBudCE4.1-DOA/DOB plasmid into T2DR4DM by nucleofection (21). The T2DR4DMDO++ [medium level of DO:DM, or (DO:DM)^M^] and T2DR4DMDO+++ [high level of DO:DM, or (DO:DM)^H^] clonal lines were constructed by fluorescence-activated cell sorting (FACS) of the parental, polyclonal T2DR4DMDO transfectant for single cells with low (e.g., 1C3) or high (e.g., 2D7) levels of surface CLIP/DR4 complexes (17). FACS-sorted single clones were then expanded in a well of a 96-well plate to establish stable, single clonal lines, including 1C3 and 2D7, as described (17). To construct T2DR4DMDO+ [low level of DO:DM, or (DO:DM)^L^], a plasmid pBudCE4.1-DOA/DOB_Gly_8_-linker_FLAG-tag was constructed similarly to the construction of pBudCE4.1-DOA/DOB. After transfection of T2DR4DM with the plasmid by nucleofection, two independent human promoters in the plasmid drove the expression of DOα and DOβ with a C-terminal FLAG-tag, respectively. Amaxa nucleofector kit C (Lonza, Basel, Switzerland) designed for nucleofection of T2 cells was used. Transfected cells were cultured in complete IMDM (IMDM-GlutaMax, 10% HI FBS and 1% P/S) with 50-100 μg/ml zeocin for 3-5 weeks to eliminate parent non-transfected cells and to construct a stable cell line, T2DR4DMDO-G8. To construct the DOKO cell line, we used the CRISPR gene-editing strategy (31) to knock out DO in T2DR4DMDO-2D7 [(DO:DM)^H^] cells. Two single guided RNA (sgRNA) primers, GGCCACCAAGGCTGACCACATGG flanking the DOA exon 1/2 and GGGGAGAAAAGTGCAACCAGAGG flanking the DOB exon 2/3 were designed using the CRISPOR website (http://crispor.tefor.net/and) and synthesized from Synthego. The Cas9 encoding plasmid was a gift from Dr. Matthew Porteus at Stanford University. sgRNA and Cas9 were introduced into T2DR4DMDO-2D7 by nucleofection using Amaxa nucleofector kit C. The nucleofection condition: 10^6^ cells in 100 μl total nucleofection solution with 8 μg of each of the two sgRNA primers and 4 μg of Cas9-encoding plasmid. Transfected cells were further scaled up and analyzed for DO expression by flow cytometry as previously described (17, 21) (see also Supplementary Fig 1 for DO expression levels).

### Cell culture

All cells were cultured in Iscove’s Modified Dulbecco’s Medium (IMDM)-GlutaMAX supplemented with 10% heat inactivated fetal bovine serum (HI FBS) and 1% of penicillin-streptomycin (P/S). All media, supplements, and selection reagents are purchased from Thermo Fisher Scientific (Waltham, MA). The expression of DR4, DM or DO in these T2 transfectants was enforced and maintained by selection with 1 mg/ml G418 (Geneticin), 1 μg/ml puromycin, or 100 μg/ ml zeocin, respectively. All cell cultures in this study were maintained in a 37 °C incubator constantly supplied with 5% CO2. Cells were harvested by centrifugation for immediate flow cytometric analysis or washed twice with ice-cold PBS and stored at −80 °C for downstream proteomic and peptidomic analyses.

### Cell line characterization using flow cytometry

To measure total protein levels by flow cytometry, T2 and its derived cell lines were fixed and permeabilized using the Cytofix/Cytoperm kit (BD Biosciences, Becton, Dickinson and Company, Franklin Lakes, NJ). Washed cells were then resuspended in 1x PermWash buffer (BD Biosciences) at a density of 1 million cells per 100 μl and stained on ice with fluorophore-conjugated mAbs. These mAbs included Fluorescein isothiocyanate (FITC)-conjugated anti-human CLIP mAb (BD Biosciences), PE (R-phycoeryhthrin)-conjugated anti-HLA-DR mAb (BD Biosciences), Alexa fluor 568-conjugated anti-human DO mAb (MagsDO5), Alexa fluor 647- or Alexa fluor 700-conjugated anti-human DM mAb (MapDM1). Alexa fluor 568-MagsDO5, Alexa fluor 647-MapDM1 and Alexa fluor 700-MapDM1 were generated previously (17, 21). To stain DR4-associated non-CLIP peptides on the cell surface, cells were pelleted by centrifugation and resuspended in phosphate buffered saline (PBS) + 1 % bovine serum albumin (BSA) at a density of 1 million cells per 100 μl, and stained on ice firstly with hybridoma supernatant containing NFLD.D11, a mouse IgM mAb known to recognize non-CLIP peptide/DR4 complexes (32), and secondly, with Alexa Fluor 647 goat anti-mouse IgM (Invitrogen, Thermo Fisher Scientific). Labeled cells were washed with corresponding staining buffer (1x PermWash buffer for intracellular staining and PBS + 1% BSA for surface staining) and resuspended in PBS + 1% BSA and analyzed on the BD LSRII flow cytometer at Stanford shared FACS Facility. Flow cytometric data were analyzed using FlowJo software (BD Biosciences).

### Protein extraction, TMT labelling and Hp-RP fractionation for proteomic analysis

Each T2-derived cell line was cultured as described above, from which three aliquots of 1 × 10^7^ cells were collected (three replicate aliquots per T2-derived cell line). To extract total protein contents for MS analysis, each replicate was lysed in 8M urea, 150 mM NaCl, 5 mM DTT, 50 mM Tris-Cl (pH 8) supplemented with Complete Protease Inhibitor Cocktail tablet (Roche, Mannheim, Germany) and 1x Halt™ Protease and Phosphatase Inhibitor Cocktail (Thermo Fisher Scientific). The lysate was then centrifuged at 13,200 rpm for 15 min, and the supernatant was transferred to a fresh test tube for a second round of centrifugation. The resulting clarified supernatant was reduced with 5 mM DTT for 30 min at 37 °C, then alkylated with 14 mM iodoacetamide for 45 minutes at room temperature in the dark and then quenched with 5 mM DTT for 20 min at room temperature. In order to clean the proteins extracted from the lysate, a methanol-chloroform precipitation was performed, and the protein pellet was washed twice with acetone. The pellet was re-suspended in 300 μl of 8M urea, 50 mM Tris-Cl (pH 8) and the concentration of total proteins extracted from a cell line was determined using the Pierce™ BCA Protein Assay Kit (Pierce, Rockford, IL).

Extracted proteins from each sample or replicate were diluted to 1 M urea, 50 mM Tris-Cl (pH 8) prior to digestion with Trypsin/Lys-C Mix (Promega, Madison, WI) at a ratio of 1:25 (enzyme: substrate; 16 hours at 37 °C). The reaction was quenched with the addition of formic acid to a final concentration of 5%. Digested peptides were desalted using a Sep-Pak C18 1 cc Vac Cartridge, 50 mg (Waters, Milford, MA). Peptides were further labeled using Tandem Mass Tag (TMT) reagents (Pierce) as previously described (33). In brief, each TMT reagent (0.8 mg per vial) was reconstituted in 40 μl of acetonitrile and incubated with the corresponding peptide sample for 1 hour. The reaction was then quenched with a final concentration of 0.3% (v/v) hydroxylamine for 15 min at room temperature. TMT-labeled peptides were acidified with 25% formic acid to pH ∼ 2. To assess labelling efficiency, a ratio-check was performed: 5 μl of TMT-labeled peptides from each cell sample or replicate were combined, desalted by StageTip (34) and then analyzed on the liquid chromatography–mass spectrometry (LC/MS)-instrument (detailed methods described below). Based on the result from the ratio-check, equal amounts of each individually labeled sample were then combined into a master pool to deliver a comparable average signal across all three TMT-labeling sets (see Fig. 1C).

**Figure 1.**
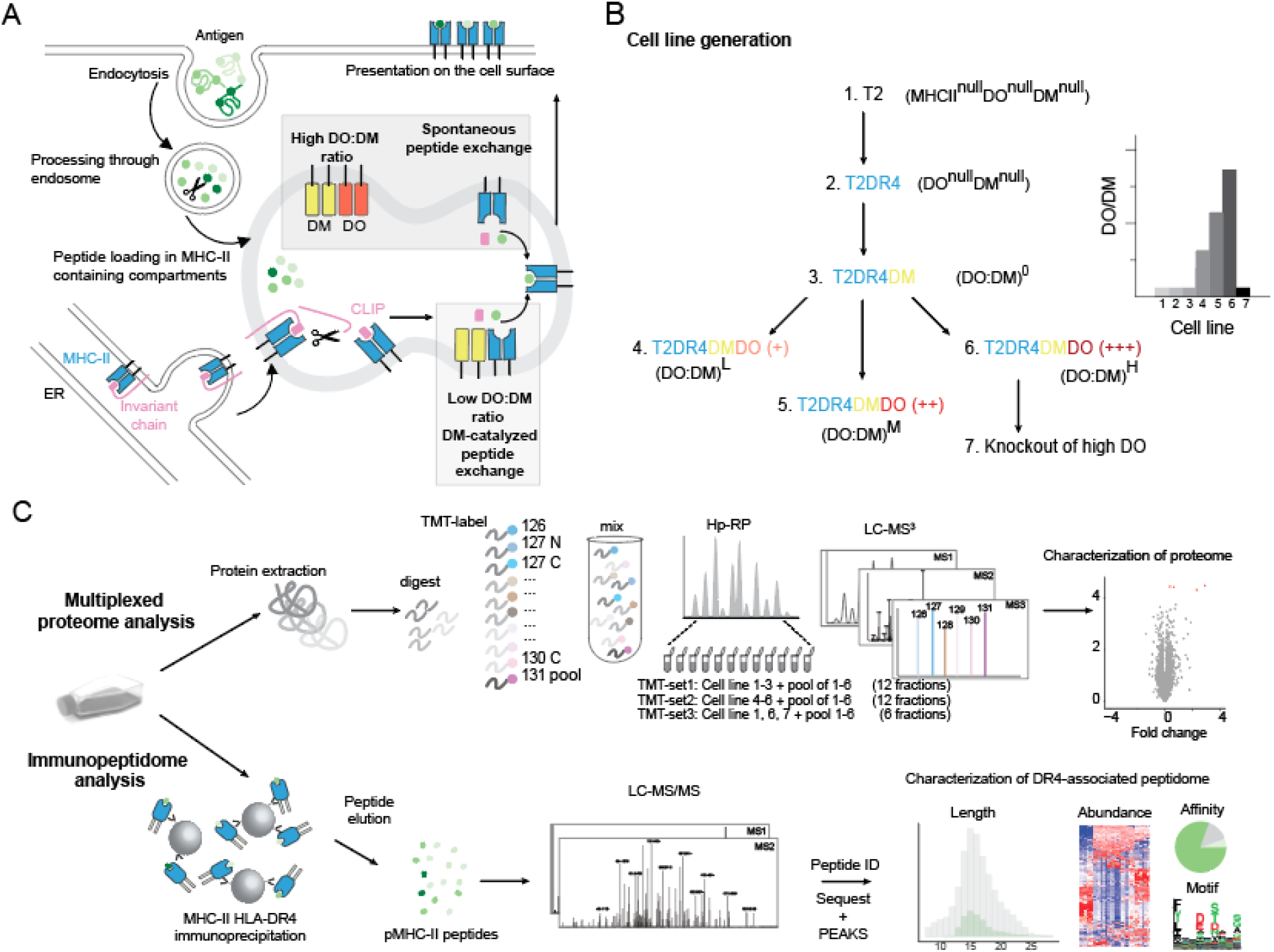
Schematic of DO-involved antigen presentation and rational for experimental design. **(A)** Model to be tested: DO association limits the abundance of free DM that catalyzes loading of peptides derived from endocytosed antigen onto nascent MHC-II thus influences the MHC-II peptidome. CLIP: class II associated invariant peptide. **(B)** Experimental design: T2-derived model cell lines expressing single MHC-II allele HLA-DR4 and different ratios of DO to DM (DO:DM) were constructed. (DO:DM)^0^, (DO:DM)^L^, (DO:DM)^M^ and (DO:DM)^H^ refer to the four DM+ cell lines with zero, low, medium and high DO:DM. (**C**) Cell lines constructed in (B) were compared with their parental cell lines by MS-facilitated multiplexed quantitative proteomic analyses and label-free peptidomic analyses of ligands eluted from MHC-II (HLA-DR4) with respect to their identities, lengths, abundances, and predicted binding properties.

TMT-labeled peptides were further desalted using a Sep-Pak C18 1 cc Vac Cartridge, 50 mg (Waters), resuspended in 10 mM ammonium formate (pH 10), and fractionated using the high-pH reverse phase (Hp-RP) fractionation approach as previously described (35, 36). For two of the TMT labeling sets, fractionation was performed using a 65 min + 15 min step-gradient buffer A (10 mM ammonium formate, pH 10) and buffer B (10 mM ammonium formate, 90% ACN, 10% H_2_O, pH 10) using an Agilent 1200 HPLC (Agilent Technologies, Santa Clara, USA). In total, 84 fractions were collected, concatenated and combined (36) into a total of 12 fractions and they were dried down. All fractions were desalted using a C18 based StageTip, dried down and stored at −80°C until final LC-MS/MS measurement. In the case of the third TMT-labeling set (containing the DOKO line) a manual Hp-RP fractionation procedure was performed using a C18 based StageTip: A total of 7 fractions were collected with sequential increases in [ammonium formate, pH = 10] concentration. The first and last fractions were combined into one fraction, while the remaining fractions remained distinct. This resulted in six final fractions. All fractions were desalted using a C18 based StageTip, dried down and stored at −80°C until final LC-MS/MS measurement.

### Mass spectrometry-facilitated proteomic analysis

Proteome-wide MS analyses were carried out as follows: TMT-labeled and Hp-RP-fractionated peptides described above were resuspended in 20 μl 0.1% formic acid, of which 10 % of the material was injected onto the Dionex Ultimate 3000 autosampler and LC-system (Thermo Fisher Scientific). Peptides were separated on a 20 cm reversed phase column (100 μm inner diameter, packed in-house with ReproSil-Pur C18-AQ 3.0 m resin by Dr. Maisch GmbH) over 120 min using a four-step linear gradient: 97% A (and 3% B) to 96% A for 15 min, to 75% A for 135 min, to 55% A for 15 min, and then to 5% A for 15 min (buffer A is 0.1% formic acid in water and buffer B is 0.1% formic acid in acetonitrile). Mass spectra acquisition was performed in a data-dependent mode on a Lumos Orbitrap mass spectrometer (Thermo Fischer Scientific, Bremen, Germany) with precursor (MS1) scans acquired in the Orbitrap mass analyzer with a resolution of 120,000 and m/z scan range 400–1,500. The automatic gain control (AGC) targets were 4 × 10^5^, and the maximum injection time was 50 ms. The most intense ions were then selected in top speed mode for sequencing using collision-induced dissociation (CID) and the fragments were analyzed in the ion trap. The normalized collision energy for CID was 35% at 0.25 activation Q. For tandem mass (MS2) scans, the AGC targets were 1 × 10^4^ and the maximum injection time was 30 ms. Monoisotopic precursor selection and charge state rejection were enabled. Singly charged ion species and ions with no unassigned charge states were excluded from MS2 analysis. Ions within ±10 ppm m/z window around ions selected for MS2 were excluded from further selection for fragmentation for 90 s. Following each MS2 analysis, five most intense fragment ions were selected simultaneously for higher energy collisional dissociation (HCD) MS3 analysis with isolation width of 1.2 m/z, normalized collision energy of 65 % at resolution of 60,000, AGC target were 1×10^5^ and maximum injection time of 90 ms.

### Computational interpretation of proteomic data

Mass spectra were initially interpreted with Proteome Discover v2.1 (Thermo Fischer Scientific, San Jose, CA). The parent mass error tolerance was set to 20 ppm and the fragment mass error tolerance to 0.6 Da. Strict trypsin specificity was required allowing for up to two missed cleavages. Carbamidomethylation of cysteine (+57.021 Da), TMT-labeled N-terminus and lysine (+229.163) were set as static modifications. Methionine oxidation (+15.995), phosphorylation (+79.966) on serine, tyrosine and threonine, and N-terminal acetylation (+42.011), were set as variable modifications. The minimum required peptide length was set to seven amino acids. Spectra were queried against a “target-decoy” protein sequence database consisting of human proteins (downloaded from the Uniprot resource, June, 2016), common contaminants, and reversed decoys of the above (37) using the SEQUEST algorithm (38). The Percolator algorithm (39) was used to estimate and remove false positive identifications to achieve a strict false discovery rate of 1% at both peptide and protein levels. Known false positives (i.e., decoys) and contaminants were excluded from further analysis steps. Differentially expressed proteins were identified, statistically evaluated, and visualized from log-transformed quantitative data using R, Rstudio, and Qlucore Omics Explorer v3.2 (Qlucore AB, Lund, Sweden). Unless otherwise noted in the text, differentially expressed proteins were selected based on a q-value threshold < 0.05 following correction for multiple hypothesis testing was applied using Benjamini-Hochberg (40).

### Purification of HLA-DR4 associated peptides

HLA-DR4 proteins were immunoprecipitated from T2-derived cells and their associated peptide cargo was eluted as previously described (41, 42), with some modifications. Briefly, each cell line was grown as two independent biological replicates, independent of cells prepared for full proteome analysis described above, and 2 × 10^8^ cells per replicate were harvested. Cells were then lysed for 20 min on ice in 20 mM Tris-HCl (pH 8), 150 mM NaCl, 1 % (w/v) CHAPS, 0.2 mM PMSF, supplemented with 1x Halt™ Protease and Phosphatase Inhibitor Cocktail (Thermo Fisher Scientific) and Complete Protease Inhibitor Cocktail (Roche). The lysate was centrifuged (2 × 30 min, 13,200 rpm at 4°C) and the resulting supernatant was precleared for 30 min using rProtein A Sepharose fast-flow beads (GE Healthcare). The precleared lysate was incubated with HLA-DR specific antibody L243 (43) (produced and purified by Genentech from hybridoma) coupled to rProtein A Sepharose fast-flow beads for 5h at 4°C. Following the immune-captures of DR4, the beads were washed with TBS (pH 7.4) and peptides were eluted from the purified DR4 molecules using 10% acetic acid. Eluted peptides were passed through a 10 kDa MWCO filter, followed by a concentration step using vacuum centrifugation, before being desalted on C18 based StageTips and stored at −80°C until LC-MS/MS analysis.

### Mass spectrometry-facilitated analysis of DR4 peptidome

Peptides eluted from DR4 proteins as described above were reconstituted in 12 μl 0.1 % FA and analyzed on an LTQ Orbitrap Elite mass spectrometer (Thermo Fischer Scientific). Samples were then injected onto the Eksigent ekspert nanoLC-425 system (SCIEX, Framingham, USA) and peptides were separated by capillary reverse phase chromatography on a 20 cm reversed phase column (100 μm inner diameter, packed in-house with ReproSil-Pur C18-AQ 3.0 m resin by Dr. Maisch GmbH) over a total run time of 160 min, including a two-step linear gradient with 4–25 % buffer B (0.2% (v/v) formic acid, 5% DMSO, and 94.8% (v/v) acetonitrile) for 120 min followed by 25-40 % buffer B for 30 min. Each cell condition was prepared in parallel as two biological replicates and each biological replicate was injected three times, each using a different complementary instrumental method, resulting in 6 raw data files per cell condition. The three instrumental methods are higher energy collisional dissociation (HCD), collision induced dissociation (CID) including single-charged species, and CID excluding single-charged species (41). Acquisition was executed in data-dependent mode with full MS scans acquired in the Orbitrap mass analyzer with a resolution of 60,000 (FWHM) and a m/z scan range 340-1600. The top ten most intense ions with masses ranging from 700-2750 Da were then selected for fragmentation then measured in the Orbitrap mass analyzer at a resolution of 15,000 (FWHM). The ions were fragmented with a normalized collision energy of 35% and an activation time of 5 ms for CID and 30 ms for HCD. Dynamic exclusion was enabled with repeat count of 2, repeat duration of 30s and exclusion duration of 30s. The minimal signal threshold was set to 500 counts.

### Identification and quantification of DR4 binding peptides from mass spectra

All tandem mass spectra were queried against the same “target-decoy” sequence database described above for the proteome analysis. All spectra were searched using both SEQUEST (38) and PEAKS DB (Studio 8, Bioinformatics Solutions Inc) (44) search engines. The MSConvert program (version 3.0.45) was used to generate peak lists from the RAW data files, and spectra were then assigned to peptides using the SEQUEST (version 28.12) algorithm. Spectra were also interpreted by the PEAKS algorithm’s de novo sequencing function to improve peptide identification confidence (41). The parent mass error tolerance was set to +/-10 ppm and the fragment mass error tolerance to 0.02 Da. Enzyme specificity was set to none and oxidation (M), deamidation (N,Q), cysteinylation (C), and phosphorylation (S, T, Y) were considered as variable modifications. High-confidence peptide identifications were selected at a 1% false discovery rate with a version of the Percolator algorithm (39) which we modified for immunopeptide analysis (41). Unlike conventional proteome analysis, false discovery rates were not evaluated at the level of assembled proteins, as this would unnecessarily penalize proteins identified by just one peptide. Quantitative abundance values (MS1 peak areas) were extracted from raw data as previously described (45). All peptide data and mass spec raw data files have been deposited in the PRIDE Archive at www.ebi.ac.uk/pride/archive (46) under accession number PXD024392.

### Characterization of DR4 binding cores and prediction of binding affinity

Immunopeptidome datasets were evaluated with the PLAtEAU script (47) to identify DR4 binding cores. Only peptides reported in both biological replicates (with the criteria of at least observed in one of the three technical replicate injections) were considered for further PLAtEAU analysis. The minimum core length was set to 13 residues (default) and the immunopeptidome data from each T2-derived cell line were analyzed individually as well as all lines and data together. The default option, “impute with lowest measured value in each run” was enabled. The quantitative cut-off criteria for being reported as differentially presented were: a p-value of less than 0.01 (corrected for multiple hypothesis testing using Benjamini-Hochberg (40)) and a log2 fold change either >2 or <-2. Both the entire set of peptides from the peptidome analysis and the subsequently defined core output from PLAtEAU were used as input for affinity prediction using NetMHCIIpan version 4.0 (48, 49). NetMHCIIpan-4.0 scores how well a given peptide sequence can bind a HLA-DR allele in question (e.g., DRB1*04:01). Its scoring models apply Artificial Neural Networks (ANNs) trained on multiple extensive datasets that measured in vitro binding affinity (BA), and MS-derived eluted ligands (EL). Optionally, NetMHCIIpan-4.0 can score binding against a model trained only on BA and not EL data. As MS data were drawn from untargeted MS studies rather than from discrete binding measurements, the two models should be expected to produce different results in some cases. We therefore designate the former prediction approach “EL”, and the latter prediction approach “BA”. For both EL and BA prediction approaches, we applied rank-based thresholds of 2% and 10% to separate strong, weak and non-binders. See main texts and figures for details. To evaluate prevalent motifs among peptides in a more untargeted fashion we applied Gibbs cluster analyses (GibbsCluster-2.0 Server (50) with the MHC-class II parameter settings) followed by visualization by Seq2Logo.

### Experimental Design and Statistical Rationale

This study was designed to include proteomic and immunopeptidomic components, which differed in the numbers of replicates and data collection modalities. Differences between these experiments were due to technical considerations, since cell input for immunopeptidome assays needed to be approximately 100-fold more than proteomic assays. Proteomic data were measured as preparative triplicates, such that each cell culture was divided into three aliquots, and each was lysed, digested, and labeled with TMT reagents in parallel as described above. Within each replicate set, all samples were processed in a single batch. LC-MS/MS analysis of HPRP-fractionated peptides proceeded according to the sequential order of the concatenated fractions, as this was not deemed to be a significant source of potentially confounding variation. LC-MS analyses were collected as single injections.

Immunopeptidomic data were collected from biological duplicates, which were processed from lysis, immunoprecipitation, and desalting at different times and in a randomized order. LC-MS analyses proceeded in a sequential order based on the nomenclature of the cell lines, one replicate set at a time. Technical triplicate LC-MS analyses were acquired per sample to increase coverage and overall signal using different experimental methods as described above. Carry-over between different samples was minimized by acquiring blank analyses between each.

## RESULTS

### Strategy for investigating the influence DO:DM has on pMHC-II repertoires

Typically, antigens undergo endocytosis and intracellular processing by specialized APCs (e.g., B cells) prior to pMHC-II presentation for CD4+ T cell scanning (7). Necessary steps in this process include proteolysis of the MHC-II chaperone invariant chain (Ii) to generate CLIP (Class II associated Ii peptide) that remains in the binding cleft of MHC-II and the spontaneous or DM-catalyzed CLIP removal and antigenic peptide loading (Fig. 1A). In B cells and several other APCs described above, this process is further modulated by DO, likely via the competition between DO and MHC-II for binding to DM (19). As only certain APC types express DO (8) and DO knockout models have yielded apparently contradictory immune consequences (11–15), the contribution of DO is puzzling. DO influences pMHC-II repertoires by controlling the level of free, active DM (17, 21, 24). High DO levels, as observed in naïve or memory B cells, limit free DM activity and permit loading of lower-affinity peptides (Fig. 1A). Acid-driven DO denaturation from DMDO complexes elevates free DM levels, thereby promoting the formation of stable pMHC-II (21). For instance, when the B cell receptor (BCR) binds antigen, B cell activation leads to acidification of late endosomes and denatures DO (17, 51) This process is BCR-antigen affinity dependent (17).

In this study, we sought to establish a tractable system to explore the peptide repertoires under varied DO:DM conditions and to elucidate the impact of DO:DM on MHC-II peptidomic landscapes. We used a series of human TxB hybrid cell lines (T2 derivatives, see Methods), each differing by only one type of protein i.e., HLA-DR4, DM and DO, or by DO:DM levels (Fig. 1B). To generate the cell lines with varied DO:DM, we used a polyclonal DO transfectant of T2.DR4.DM cells and two clonal lines expanded from FACS-sorted single cells, based on levels of surface CLIP/DR4 complexes (Fig. 1B, see also Methods). We extracted all protein content from the full panel of cell lines for mass spectrometry (MS)-facilitated proteomic analysis; in parallel, we eluted DR4-associated peptides for MS peptidomic analysis (Fig. 1C). This strategy allowed us to measure how DO:DM influences DR4 peptidomes, while accounting for the role protein abundance may have in peptide selection, and any subtle proteome-wide alterations that might exist between each cell line.

### pMHC-II-specific mAbs reflect DO:DM influence on DR4 peptide presentation

We assessed the expression levels of MHC-II and accessory molecules in the full panel of T2-derived stable cell lines. MS-facilitated proteomic analysis validated the expression of α and β chains of DR, DM, and DO (Fig. 2A, Supplementary Dataset 1). Their levels were consistent with those of the corresponding total αβ heterodimeric protein detected by flow cytometry (Fig. 2B). Flow cytometry analysis further showed that DM+ lines have comparable levels of total DR and a hierarchy of total DM or DO (Fig. 2B and Supplementary Fig. 1). The concurrent decrease of DM levels as DO levels increased was a consequence of how (DO:DM)^M^ and (DO:DM)^H^ lines were selected based on CLIP/DR4 at the cell surface. This resulted in a step-wise increase of DO:DM across these cell lines. As we observed previously (19, 21), DM expression, unopposed by DO in the (DO:DM)^0^ line, catalyzed the removal of almost all CLIP bound by DR4. In contrast, CLIP/DR4 persisted in T2DR4, which lacks DM and DO. DO expression rescued CLIP/DR4 in (DO:DM)^H^, whereas CLIP level was only slightly increased in (DO:DM)^L^ and (DO:DM)^M^ compared to (DO:DM)^0^. Using NFLD.D11, an antibody specific for non-CLIP/DR4 (32), we further determined that (DO:DM)^H^ displayed significantly lower levels of non-CLIP/DR4 than the levels in the other DM^+^ lines. The high level of invariant chain protein (CD74, see Fig. 2A) found in (DO:DM)^L^ and (DO:DM)^M^ concurrent with low levels of CLIP/DR4 (Fig. 2B) suggests that CLIP/DR4 levels do not depend on CD74 protein abundance. Instead, these data suggest CLIP/DR4 levels directly depend on DO:DM stoichiometry. Interestingly, non-CLIP/DR4 levels, as detected by NFLD.D11, were increased in (DO:DM)^L^ and (DO:DM)^M^ compared to (DO:DM)^0^, indicating the dependence of some non-CLIP peptides on expression of some DO, in addition to DM.

**Figure 2.**
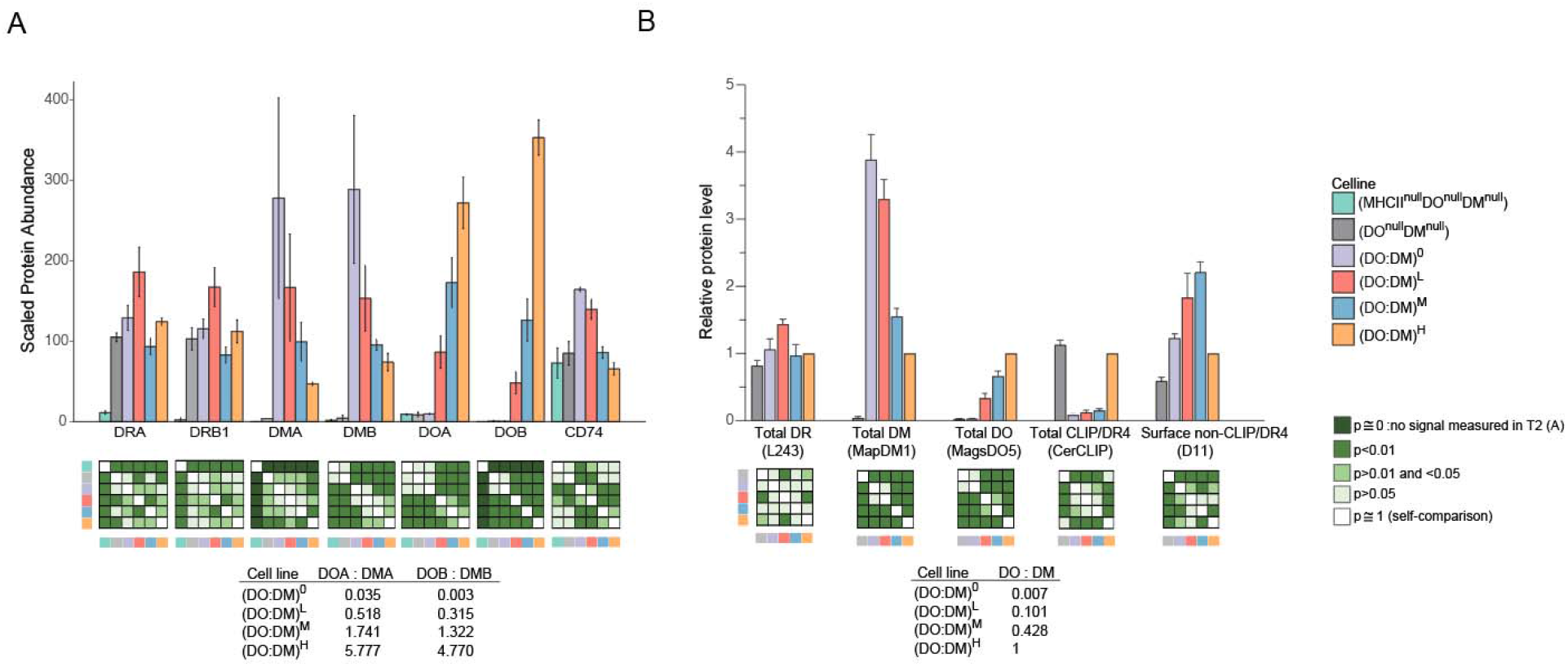
T2-derived cell lines demonstrate anticipated abundances of HLA-DR and associated chaperones or peptides. **(A)** Mass spectrometric analysis of the protein abundance for the α or β chains of DR, DM or DO, and for CD74 (the invariant chain). Data represent scaled TMT-labeled peptide signals from the corresponding protein ± SD. DMA and DOB were not detected in parental T2 (DR^null^DO^null^DM^null^) cells. **(B)** Flow cytometric analysis showing the relative protein level for total DR, DM, DO or CLIP/DR4 complexes, and the surface level of non-CLIP/DR4 complexes. Monoclonal antibodies (mAb) used for staining are L243, MapDM1, MagsDO5, CerCLIP, and NFLD.D11 as indicated. Data represent mean fluorescence intensities of the corresponding staining in each cell line normalized by the (DO:DM)^H^ line + SEM. See Supplementary Fig. 1 for representative flow histograms. In both (A) and (B), a two sample t-test was used to compare each pair of the indicated cell lines for the indicated protein level. The calculated ratios of DO, DOA, or DOB to DM, DMA or DMB, respectively, reflect the relative abundance differences rather than the absolute number in the four DM-expressing lines.

Overall, these results indicate a substantial and nuanced influence DO:DM can have on pMHC-II presentation, an effect which varies with DO:DM. Thus, the T2-derived lines described here represent the first system for directly measuring DO:DM impact on HLA class II peptidomes. Of note, DO:DM can be directly measured, whereas its functional consequence – free, active DM – cannot. Thus, we have designated cell lines by this ratio. Across the full panel of cell lines, free DM is highest in the (DO:DM)^0^ cells, and decreases as the DO:DM ratio increases. It is fully absent in the DO^null^DM^null^ cells. While this last configuration of DO, DM and DR expression is unlikely to be found under physiological conditions, it could nevertheless reflect DM-resistant MHC-II alleles (52, 53).

### DR4 peptidomic differences between cell lines corroborate synergistic DO:DM tuning

To evaluate how DO:DM affects DR4-presented peptide repertoires, we eluted peptide ligands from DR4 in each cell line and analyzed them by mass spectrometry (Supplementary Dataset 2). We identified an average of 3,072 unique peptides per biological replicate analysis per cell line, and 10,587 unique peptides from the entire data set (Supplementary Table 1, Supplementary Dataset 2). These peptides had an average length of 16.6 amino acids (aa) (Fig. 3A), an appropriate average length for MHC-II associated peptides. Peptides with no quantified signal were excluded from further consideration, resulting in 7,390 quantified unique sequences. Of these, 4,224 (4,528 if considering different residue modifications) were quantified in both bio-replicates.

**Figure 3.**
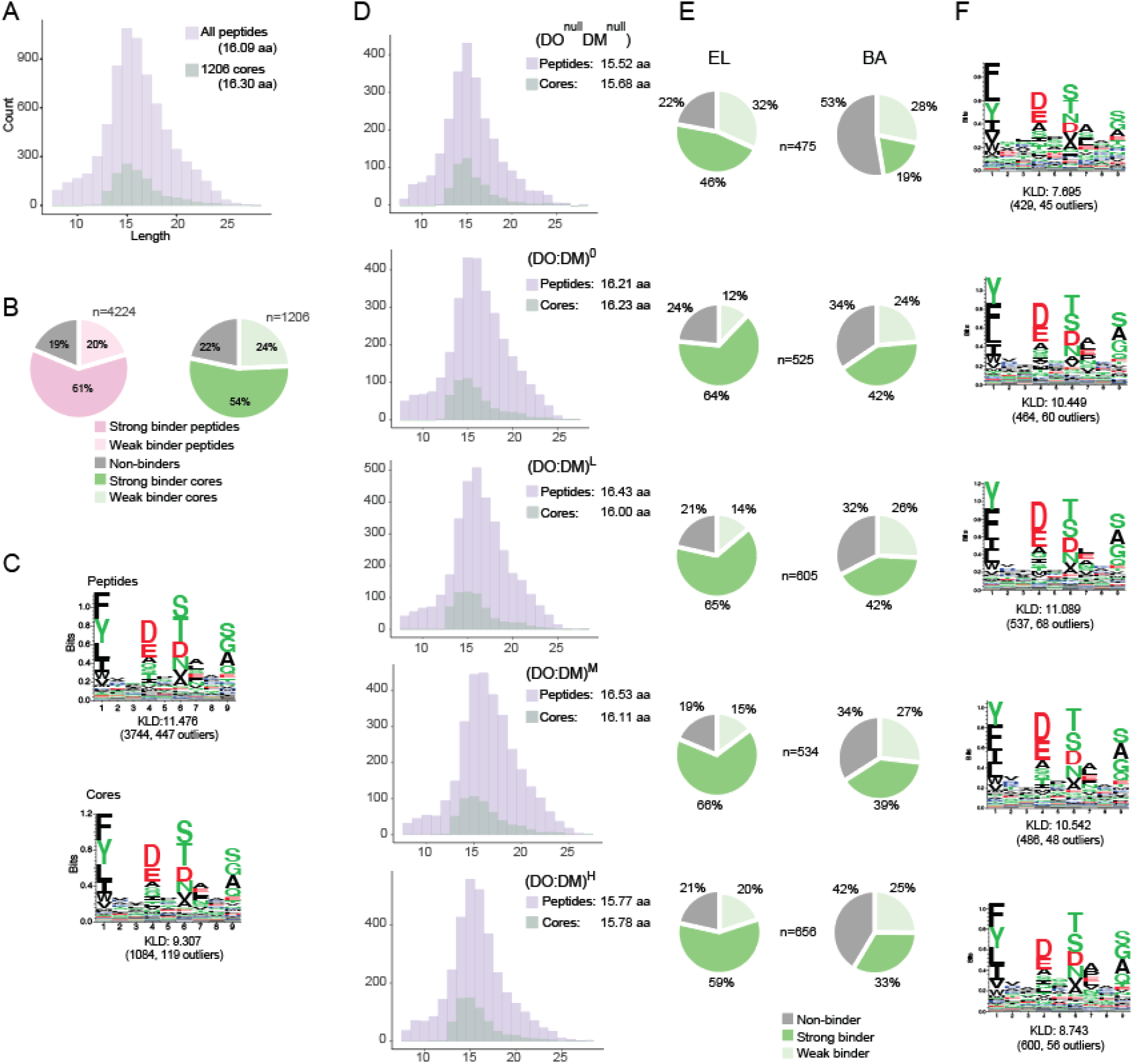
Characterization of the combined DR4 peptidome from 5 cell lines or peptidomes from each individual line. **(A)** Length distribution for MS-identified unique peptides from all 5 lines and their consensus cores determined using the PLAtEAU algorithm. The mean lengths of each are indicated in parentheses. **(B**) Percent of peptides or cores predicted to be strong (top 2%rank), weak (2-10%rank) or non- (bottom 90%rank) DR4-binders as predicted by NetMHCIIpan-4.0 (EL approach, see Methods). (**C)** DR4 motifs generated with Seq2Logo. All peptides (top) or all cores (bottom) described in (B) were first clustered with GibbsCluster. Motifs indicate frequencies of particular residues at each of the nine positions of the DR4 binding register, with increased aa specificity at anchor residue positions (P1, P4, P6, P9). The top-ranked cluster for each condition is presented. The Kullback-Leibler Distance (KLD) score is listed and the size of the cluster and number of outliers is listed in brackets. **(D-F**) Analyses of length distributions (D), percent of cores predicted to be strong or weak binders using the eluted ligand (EL) and in vitro binding affinity (BA) models implemented by NetMHCpan-4.0 (see Methods) (E), and motifs deduced from DR4-binding cores (F) identified from each individual cell line.

Most MHC-II-eluted peptides we identified could be grouped into nested sets of overlapping peptides. This is a common observation, resulting from the MHC-II open-ended binding groove that permits peptides of multiple lengths to bind (54). This peptide sequence heterogeneity can complicate subsequent computational procedures, such as sequence motif determination and ligand quantification. We implemented the PLAtEAU algorithm (47), which addresses this challenge by condensing overlapping peptides to single DR4-binding peptide cores. These cores represent consensus sequences shared by nested sets of peptides, each of which should contain the same binding register with the expected P1, P4, P6 and P9 anchor positions that directly interface with the MHC-II binding groove. Applying this approach effectively simplified our subsequent analyses, while aggregating analyte signals that were sometimes distributed across multiple peptides (Supplementary Figure 2). PLAtEAU determined 1,206 unique peptide binding cores with an average length of 16.30 aa (Fig. 3A, Supplementary Table 2). Each of these cores, which mapped to 782 self-proteins (Supplementary Fig. 2), contained at least one 9-aa register expected to occupy the DR4 peptide-binding groove. The NetMHCIIpan-4.0 algorithm (48, 49) predicted that 81% of these peptides and 78% of these cores ranked among the top 10% of sequences able to bind DR4 (Fig. 3B). The DR4 binding motifs generated from the entire peptidomic dataset and from the condensed cores (Fig. 3C) corresponded well with the reference motifs generated using the NetMHCIIpan-4.0 motif viewer (Supplementary Fig. 3). Collectively, the core dataset and the MS-derived peptide dataset from which it is derived, both cover informative DR4-presented peptides with expected lengths, motifs, and predicted binding features, and both are therefore appropriate for further analysis.

We next compared the condensed core pMHC-II repertoires derived from each individual cell line. Compared to DO^null^DM^null^ or (DO:DM)^H^, DM^+^ lines with zero to medium DO:DM showed a tendency towards longer peptides or cores (Fig. 3D, Supplementary Fig. 4B). These lines, which have modest DO:DM, also yielded a greater proportion of predicted strong DR4-binding cores (Fig. 3E), and similar motifs to one another (Fig. 3F). The amino acid residue preference at the four dominant anchor positions (P1, P4, P6, P9) were most similar among the (DO:DM)^0-M^ lines, suggesting a DM-mediated preference for Y over F at P1. We note, however, that the top two most frequent residues at each of the anchor positions were conserved among all five lines (Fig. 3F). Interestingly, the (DO:DM)^H^ line generated the greatest number of cores (656), compared to (DO:DM)^0-M^ (Fig. 3E, F); this indicates that the presence of DO enables a larger number and an increased diversity of peptides to be presented by DR4, as also observed by Nanaware et al, for DR1 (30). These differences are consistent with greater free DM levels in (DO:DM)^0-M^ than (DO:DM)^H^, with consequent selection for high affinity, stable pMHC-II complexes with decreased sequence diversity. Equivalent results between peptide and binding core sequences (Supplementary Fig. 4 and 5) support the non-redundant core analysis as an appropriate data simplification tool.

### Restricted DO:DM ratios tune distinct subsets of peptides for DR4 presentation

From a global perspective, differences between each cell line were most apparent relative to DO^null^DM^null^ or comparing (DO:DM)^H^ to (DO:DM)^0-M^ (Fig. 3). Within any of these comparisons however, specific peptide cores could demonstrate a range of abundance profiles. To develop a more granular understanding of DO:DM influence on antigen presentation, we directly compared the abundances of each peptide core between all cell line pairs (Fig. 4A,B). We found smaller differences between each of the three (DO:DM)^0-M^ cell line pairings and larger differences between (DO:DM)^0-M^ and DO^null^DM^null^, as compared to the differences observed between the other pairs of cell lines (Fig. 4B, comparisons 1-3 or 8-10 versus comparisons 4-7). This analysis confirmed the greater similarity between DR4 peptidomes derived from the three (DO:DM)^0-M^ lines, as mentioned above (Fig. 2A and 3E,F). This grouping is surprising considering that (DO:DM)^L^ and (DO:DM)^M^ expressed DO whereas (DO:DM)^0^ did not. This suggests that many peptide cores did not simply increase or decrease monotonically with tuning the DO level itself; rather different peptides might be dependent on a synergistic tuning of DO:DM. Particular DO:DM thresholds might therefore promote some peptides’ efficient loading onto DR4 while inhibiting others.

**Figure 4.**
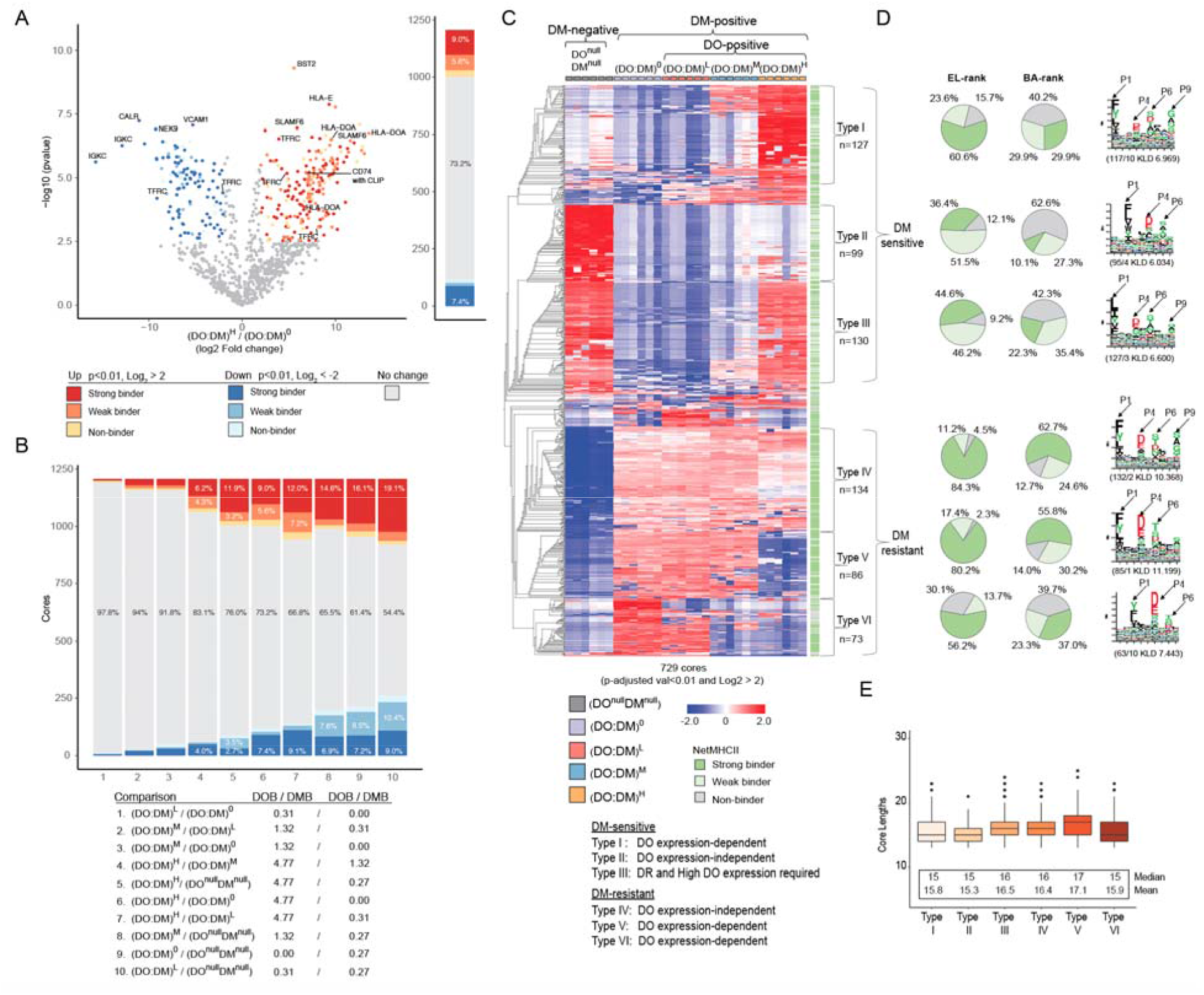
Classification of distinct binding core types. **(A)** A representative volcano plot illustrating abundance differences of cores between two cell lines, (DO:DM)^H^ and (DO:DM)^0^. Significance cutoff: p<0.01 of t-test after being adjusted for multiple hypothesis testing (BH) and a Log2 fold change >2 (in red color scale) or <-2 (in blue color scale). Strong, weak, or non-binders predicted using the NetMHCIIpan-4.0 EL model are indicated by different color intensities. The source proteins of a selected subset of cores are indicated. The stacked bar to the right aggregates the seven types of data points displayed in the volcano plot. **(B)** Summary of all pair-wise comparisons between five cell lines. Stacked bar chart based on differential presentation, similar to that described in (A). Cell line details with the MS-determined DO:DM are indicated. **(C)** Heatmap (z-score normalized and hierarchically clustered) showing cores with significantly different abundances between at least two cell lines using criteria described in panel (A) and classification of six core types. Green scales to the right indicate the corresponding core’s binding prediction using NetMHCIIpan-4.0’s EL model. The six columns per cell line represent two biological replicates, each with three technical replicate LS-MS analyses. **(D)** Percent of Type I-VI cores predicted to be strong-, weak-, or non-binders using EL or BA approaches (see Methods), and the corresponding sequence motif extracted from each, as described in Fig. 3. **(E)** Length distribution for the six types of cores.

To explore DO:DM tunable profiles further, we clustered cores based on their relative abundance across cell lines (Fig. 4C). Consistent with our finding that most (729 of 1,206, Supplementary Table 3) peptide cores were presented with significantly different abundances in at least two of the five cell lines, this clustering analysis revealed 6 major core types: Type I: [DO^null^DM^null^, (DO:DM)^0-L^] < (DO:DM)^H^; Type II: (DO:DM)^0-H^ < DO^null^DM^null^; Type III: (DO:DM)^0-M^ < [(DO:DM)^H^ and DO^null^DM^null^]; Type IV: DO^null^DM^null^ < (DO:DM)^0-H^; Type V: [DO^null^DM^null^, (DO:DM)^H^] < (DO:DM)^0-M^; Type VI: [DO^null^DM^null^, (DO:DM)^M-H^] < (DO:DM)^0-L^. The majorities of Types II and III (up to 51.5%) were predicted weak binders, whereas the majorities of Types IV and V (up to 84.3%) were predicted strong binders, regardless of prediction approaches used (Fig. 4D). The deduced motif types associated with these peptide core types were related to the canonical DR4 binding motifs (Supplementary Fig. 3).

These DR4-binding and abundance characteristics (Fig. 4C and Supplementary Fig. 2A) suggest these binding cores fall into two primary categories: DM-sensitive (Types I-III) and DM-resistant (Types IV-VI). DM-sensitive cores may spontaneously bind DR in the absence of DM notably in DO^null^DM^null^ cells. These cores may be readily displaced or competed off by DM-resistant cores in DM’s presence. This appears to be the case particularly for Types II and III, half of which are predicted to be weak binders (EL-rank, Fig. 4C, D). For others such as Types I and III, binding to DR in the presence of DM can be protected with high DO:DM, which effectively opposes when DM activity. In contrast, DM-resistant cores are presented almost only in the presence of DM, and many are predicted to be strong DR4-binders (Fig. 4C, D). Some like Type IV can be selected for DR4 presentation at higher DO:DM condition when there is less, but apparently adequate, free DM activity. In contrast, others like Type VI are not presented until DO:DM reaches very low levels.

Of note, cores of Types II and V showed the shortest and longest average core lengths, respectively (Fig. 4E), in accordance with their predominantly weak and strong binding affinities as predicted using NetMHCpan-4.0’s BA model. This trend was less apparent from the EL model’s binding predictions (Fig. 4D). The longer peptide lengths associated with significant free DM may also reflect DM-mediated peptide selection in earlier, less acidic endosomal compartments with reduced exopeptidase activity; these are compartments where DO, if present, would effectively inhibit DM. The dominant affinity group for Types I or VI also showed some discrepancies when using EL versus BA prediction approaches. Both observations reflect limitations, perhaps related to NetMHCII-4.0 prediction methods (see discussion). In addition, Types II and VI showed an unusual shift of the anticipated dominant anchors (Fig. 4D; P1 is located at the 3^rd^ position in these cores whereas it is in the 1^st^ position in the other Types). This shift, which implies longer N-terminal extensions among the underlying peptides, could reflect algorithmic limitations (see also Discussion) due to suboptimal input (<100 core peptide sequences) for motif generation.

### pMHC-II characteristics contribute to DO:DM-tuned peptide presentation

The intra-binding core Z-score normalization we used to generate Figure 4C effectively showed relative abundance changes between each cell line for a given core, but could mask gross abundance differences between the various binding cores we measured. To further examine how tuning DO:DM can affect different subsets of peptides presented by DR4, we compared the relative abundances of each peptide core type across all cell lines (Fig. 5A). Despite the similar numbers of cores clustered into each of the three DM-sensitive core types (Fig. 4C), Types I and II showed substantially lower abundances than Type III (Fig. 5A). Thus, types I and II account for a small contribution to the overall peptidome in any of the cell lines. In contrast, the abundances associated with the DM-resistant cores of types IV-VI appeared to scale approximately with the number of cores represented in each group (Fig. 5A and 4C). Of these, type IV was the most abundant peptide type in the 3 cell lines with substantial DM activity [(DO:DM)^0-M^] (Fig. 5A and 4C), but notably a strong binding Type IV subset was enriched when DM activity was most limited by DO (Fig. 4C).

**Figure 5.**
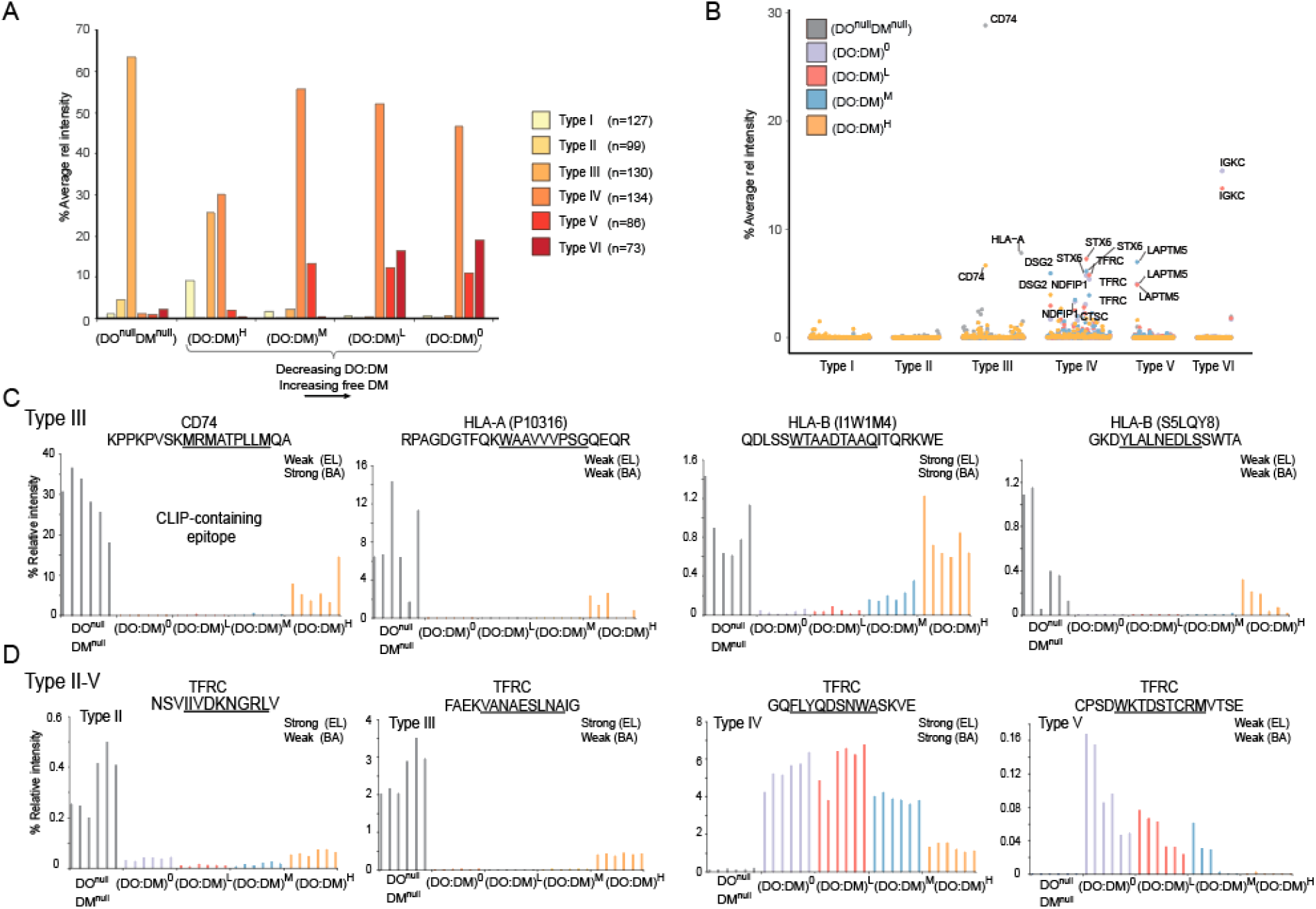
Abundance comparisons between each core type across the DM:DO gradient. **(A)** Percentage abundance subtotal of each type of cores (clustered in Fig. 4C). Percentage relative average intensity spanning the two biological replicates displayed for the six types. In addition, the residual % relative intensity covering all the six subtypes (I-V) is also displayed. **(B)** Percentage abundance subtotal of each core (clustered in Fig. 4C). **(C)** Four examples of type III cores from CD74, HLA-A and HLA-B. **(D)** Example of cores for Type II-VI for the transferrin receptor protein 1 (TFRC).

To better understand how DO:DM effects might differentially apply to specific DM-sensitive or-resistant peptides, we evaluated how cores’ abundances (Supplementary Table 2) contributed to these Type-specific observations (Supplementary Fig. 6). Among the DM-sensitive Type III core sequences, the CLIP (CD74) sequence, KPPKPVSK<u>MRMATPLLM</u>QA (binding register underlined), was dominant, with 28.8% normalized intensity in DO^null^DM^null^ and 6.66% in (DO:DM)^H^ (Fig. 5B). This core represents the most abundant overlapping peptide sequences in our entire dataset (Supplementary Dataset 2). The CLIP core demonstrated greatly reduced abundance in (DO:DM)^0-M^ cells (0.05%, 0.14% and 0.20%, respectively; Fig. 5C). The trend was consistent with the flow cytometric measurements of CLIP/DR4 (Fig. 2B), and confirms the known role DO and DM have on catalyzing peptide exchange for CLIP. The second most abundant DM-sensitive Type III core (RPAGDGTFQKWAAVVVPSGQEQR), with 7.8% normalized intensity in DO^null^DM^null^ (Fig. 5B, 5C) was derived from the HLA-A class I protein. It also followed a similar decrease in abundance (normalized intensities of 0.02%, 0.01%, 0.03%) across the (DO:DM^0-M^ gradient) with a substantial increase in (DO:DM)^H^ (1.2%, Fig. 5C). Two HLA-B-derived cores demonstrated similar DM-sensitive profiles across the five cell lines, albeit with ten-fold lower intensities (Fig. 5C). A majority of DO:DM-dependent reductions in peptide abundance did not reflect any underlying protein abundance differences and proteins were consistent across the five cell lines (Supplementary Fig. 7). Exceptions were expected, such as a core derived from HLA-DOA, which was undetectable in the DO^null^DM^null^ cell line. Overall, these results support the notion that DM selects against subsets of weak-binding peptides in a DO-dependent fashion.

In Type III DM-sensitive cores, large overall abundance swings (Fig. 5A) were dominated by a small number of cores (e.g., CLIP/CD74) in DO^null^DM^null^ and (DO:DM)^H^ cells (Fig. 5B). In contrast, the greater abundance of DM-resistant cores of Types IV-VI among (DO:DM)^0-M^ cells (Fig. 5A) can be attributed to a more diverse set of cores, each with moderately high abundance – for example, binding cores derived from DSG2, IGKC, LAPTM5, NDFIP1, STX6, and TFRC (Fig. 5B). These were mostly concentrated among Type IV cores, whereas types V and VI core abundances were more restricted to single protein sources (LAPTM5 and IGKC, respectively). As with the DM-resistant cores, these patterns relative to DO^null^DM^null^ cells cannot simply be attributed to abundance differences at the protein level (Supplementary Fig. 7). Instead, these three DM-resistant core types appear to have distinct DO:DM thresholds that govern their presentation by DR. We attributed this differential peptide presentation to the graded DO:DM effects on DM selection for high affinity, stable peptide/DR4 complexes, as most related cores were predicted by NetMHCIIpan-4.0 to be strong binders.

These data further demonstrate that single proteins could harbor cores of multiple Types. For example, we identified multiple cores across the length of TFRC with moderate to high abundance belonging to four core types (II-V) (Fig. 5D): In agreement with the aggregated quantifications in Fig. 5A of DM-sensitive peptides, the Type III peptide FAEKVANAESNAIG was approximately ten-fold more abundant than the Type II peptide NSVIIVDKNGRLV in DO^null^DM^null^ and (DO:DM)^H^ cells, but neither were robustly presented in (DO:DM)^0-M^ (Fig. 5D). In contrast, the Type IV and Type V DM-resistant cores, GQFLYQDSNWASKVE and CPSDWKTDSTCRMVTSE, decreased to different extents with increasing (DO:DM)^0-H^ (Fig. 5D). Of these peptides, the most abundant (DM-resistant Type IV, GQFLYQDSNWASKVE, up to 6% relative intensity in (DM:DO)^0-L^) was predicted to be a strong binder by both EL and BA models (Fig. 5D). The other DM-resistant peptide (Type V, CPSDWKTDSTCRMVTSE) had more than 30-fold lower intensity in (DM:DO)^0-H^ than the above peptide. In agreement with this abundance difference, it was predicted to be a weak binder by both EL and BA models (Fig. 5D). In contrast, both DM-sensitive peptides above were predicted to be strong binders by the EL model, and weak binders by the BA model, despite differing by approximately 10-fold in their abundances. These observations underscore the notion that tuning DO:DM ratios can asynchronously shift the abundances of cores derived from a particular protein.

### DO knock-down supports DO:DM tuning of DR4-presented peptides

DO is downregulated in naïve and memory B cells that enter GC for antigen presentation and affinity maturation of their antigen receptors. To mimic this regulation and to verify our model, we used CRISPR (31) to knock down DO levels in (DO:DM)^H^ cells (DOKO) and measured the corresponding DR4 peptidomes and proteomes using the aforementioned strategy (Figure 1C, Supplementary Fig. 8). Flow cytometric assays indicated lower levels of the DO heterodimer in the DOKO cell line compared to (DO:DM)^H^ cells (Fig. 6A, Supplementary Fig. 8). This was confirmed by MS-based proteomic measurements of the DO α and β chains (Fig. 6A). Notably, using CRISPR to edit the DO gene preserved a similar degree of cell heterogeneity for DO expression (Supplementary Fig. 9). This should also reflect a similar scope of DO downregulation as might be observed in physiologic contexts. We further confirmed that 99.9% (4,060 of 4,064) quantified proteins (TMT-label set 3) demonstrated consistent expression between the DOKO and (DO:DM)^H^ cell lines (Fig. 6B). Notable exceptions included the DO α and β chains (both p<0.01 and log2 fold change >2) that we specifically targeted by CRISPR.

**Figure 6.**
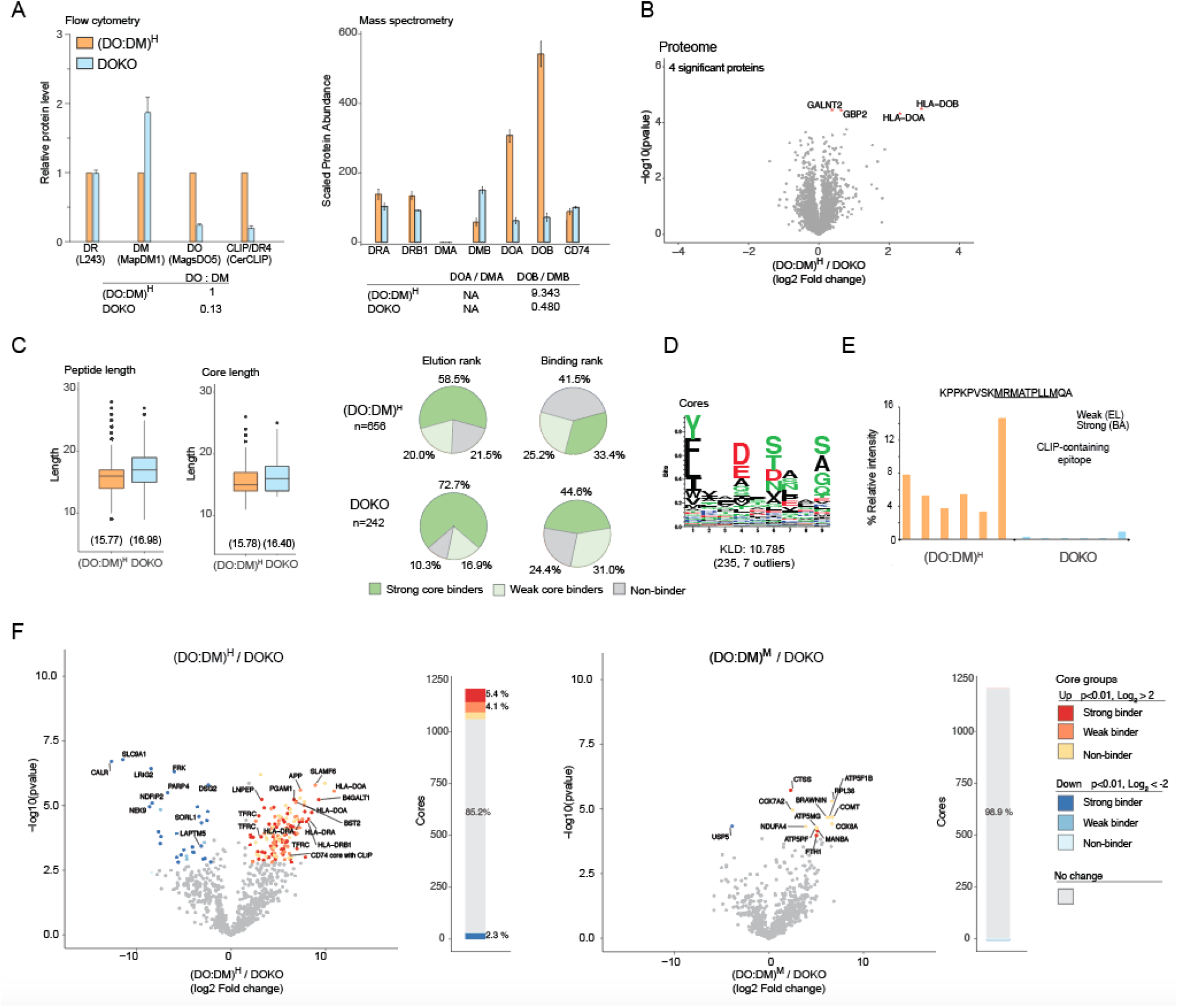
CRISPR-mediated transformation of high DO:DM to low replicates (DM:DO)^L^ cell phenotype. (**A**) Flow cytometric analysis and mass spectrometry quantification of DR, DM, DO and CLIP. Relative protein expression levels between the cell lines for DR, DM, DO and CD74 based on total cell lysate mass spectrometry-based TMT-proteomics data. No peptides were sequenced from the DMα (DMA) gene in this TMT-labeling set, resulting in undefined ratios involving this protein. **(B)** Minimal differential proteome expression differences were observed between the (DO:DM)^H^ line and its knockout. Differentially upregulated expressed proteins with a q-value of <0.05 in the (DO:DM)^H^ line are indicated in red. **(C)** Peptide length, core epitope length and binding prediction. **(D)** DOKO motif generated as in Fig. 2C. The Kullback-Leibler Distance (KLD) score is listed and the size of the cluster and number of outliers is listed in brackets. (**E)** DOKO demonstrated markedly reduced levels of the dominant CLIP containing core. **(F)** Differential presentation between DOKO cells and its parental (DO:DM)^H^ line or the lower DO:DM cell line (DO:DM)^M^. As in Figure 4A, data are presented both as volcano plots and stacked bar charts based on differential presentation (t-test, with a p<0.01 after being adjusted for multiple hypothesis testing (BH) and a Log_2_ fold change larger than 2 or smaller than −2. Cores with different binding affinities (predicted by NetMHCIIpan-4.0) using elution rank are indicated by different color intensities. Proteins which gave rise to a selected subset of relevant cores are indicated.

Consistent with the aforementioned difference between cores presented in (DO:DM)^H^ and (DO:DM)^0-M^, increased free DM in the DOKO resulted in longer peptides and longer cores, a larger proportion of which were predicted strong binders (Fig. 6C). The sequence motif generated from the DOKO peptidomic data (Fig. 6D) was more homologous to that generated using the (DO:DM)^0-M^ peptidomic data (Fig. 3F) than either DO^null^DM^null^ or (DO:DM)^H^ cells, consistent with free DM activity. As expected, a drastic reduction in the dominant CLIP containing core was observed in the knockout. (Fig. 6E). In keeping with their lower DO:DM ratio, pair-wise volcano plot comparisons indicated that DR4-presented peptides in DOKO cells were qualitatively more similar to those presented in (DO:DM)^M^ cells than (DO:DM)^H^ cells (Fig. 6F), consistent with the overall expected peptidomic outcome after DO:DM was reduced (Supplementary Fig. 9A). Considering all cores mentioned earlier (Fig. 5C,D), abundances in DOKO changed from the level in (DO:DM)^H^ to a level that was similar to (DO:DM)^0-M^ (Supplementary Fig. 9B,C, Supplementary Fig. 10). These changes were dependent on decreased DO expression and the corresponding tuning of DO:DM, not the differential expression of the proteins from which these cores were derived (Supplementary Fig. 7). Collectively, the knock-down study demonstrates that a major function of DO expression in specialized APCs is to maintain an appropriate DO:DM ratio, the tuning of which controls DM editing and the resulting DR4-presented peptide repertoires.

## DISCUSSION

In previous studies, we measured correlations between DO:DM and DM-catalyzed peptide exchange using soluble proteins (21) B cell lines, and *ex vivo* human tonsillar B cells (17) as three model systems. Our findings suggested that the tunable DO:DM is a crucial factor governing free DM levels and the consequent spectrum of presented peptides. This kind of regulation could help explain how different APCs expressing the same complement of MHC-II alleles and exposed to the same antigens, might present different peptides in different contexts: as naïve and memory B cells enter the GC, and as certain DC mature, they downregulate DO from otherwise high expression levels. Accordingly, these APCs can experience widely ranging DO:DM states, and consequently, pMHC-II repertoires bearing markedly different DM editing signatures. If one were to simply consider DO expression as either on or off, many antigen-specific outcomes – particularly when measured as T-dependent immune responses – would not be explained (11, 14, 24–30).

Considering the limitations of only examining two DO expression states (DO+ versus DO-), we developed a model system to reevaluate DO regulation by creating three DO+ states and two DO-states, covering “infinitely” opposed (No DM function in DO^null^DM^null^ cells), high, medium, low and zero levels of DO:DM. MS-facilitated peptidomic and proteomic measurements demonstrated the critical tuning function DO:DM can have in modulating HLA-DR4-presented peptidomes. Evidence from DO knockdown relative to (DO:DM)^H^ cells (highest DO level) supports the idea that DO downregulation could create multiple DO:DM states. Although each DO:DM cell state yielded a different spectrum of peptides with restricted DM sensitivity ranges, we found that decreasing DO:DM increasingly favored peptides predicted to be DM-resistant. This trend predicts that upon antigen exposure, DO downregulation in DO-expressing APCs would promote increased loading of DM-resistant antigenic peptides onto DR.

From many self or foreign peptides with the potential to bind DR with high affinity, those that survive DO:DM-tuned DM editing can be selected based on multiple factors. These include the abundance of source proteins and their sequence-susceptibility to proteolysis; inherent allelic characteristics and intracellular modifications of MHC-II that influences binding specificities; intracellular modifications of source proteins or their peptide derivatives; and endolysosomal locations where proteins are degraded or protected in a pH-dependent fashion. Our MS-based peptidomic and proteomic analyses across seven DO:DM cell states reveals several pMHC-II presentation characteristics relate to these factors in addition to DO:DM tuning, as discussed below.

First, distinct DO:DM thresholds correspond to different free DM activity levels, and therefore the different types of peptides that survive DM editing. Although DO can block DM-catalyzed CLIP removal from DR4 (55), our model demonstrated that CLIP can be effectively replaced by other peptides when DO:DM is reduced to moderate levels (i.e., (DO:DM)^M^ or even (DO:DM)^H^, Fig. 4,5). For example, Type IV peptides with high predicted binding affinity (both BA and EL models) like TFRC GQFLYQDSNWASKVE (Fig. 5D, Supplementary Fig. 10) were presented by all cells with any DM activity, including (DO:DM)^H^. In contrast, other peptides with low predicted binding affinities (BA model) were either included (TFRC FAEKVANAESLNAIG) or excluded (TFRC CPSDWKTDSTCRMVTSE) among peptides presented by (DO:DM)^H^ cells. Overall, nuanced DO:DM specificity patterns were apparent from our data set (Fig. 4C). In addition, it seems likely that an allelic specificity component to determining the DO:DM thresholds or free DM levels that effectively evacuate the peptide-binding groove to accommodate new peptides. To an extreme extent, some HLA-DQ2 alleles have evolved to require very high DM expression to overcome their intrinsically low DM-susceptibility (52). The DRB1*04:01 allele studied in our model has lower affinity for CLIP than many other MHC-II alleles (56). As a result, spontaneous exchange of other peptides was observed across all DO:DM states in these cells.

Second, in addition to the underlying DO:DM context, a peptide’s DM-sensitivity, binding affinity and abundance influence the likelihood it will contribute to pMHC-II repertoires. Although our assessment of binding strength was based on scoring with predictive algorithms (BA and EL models) rather than direct measurement, it is likely true that eluted peptides from each DO:DM cell state and across all six types of cores include both strong and weak binders. However, a large proportion of Types I-III cores from DO^null^DM^null^ and (DO:DM)^H^ cells were predicted to be weak binders and were more abundant (particularly CLIP peptides from CD74 and HLA class I) than strong predicted binders from the same core Type. A large proportion of cores among DM-resistant Types IV-VI from (DO:DM)^0-M^ were strong binders (e.g., cores derived from STX6, LAPTM5, and TFRC (Fig 5) and had relatively higher abundance than weak binders from the same categories. Therefore, we propose that APCs are regulated by DO:DM tuning to present substantial amounts of peptides with binding strengths reflecting the degree of DM editing, while allowing peptides with wider affinity ranges to be presented at lower abundances. One caveat is that our analysis was based on isolating total DR4 rather than surface-expressed molecules; thus in (DO:DM)^0-M^ cells, pMHC-II with weaker binding peptides might represent intermediates along the presentation pathway.

An implication of this model is that when building MHC-II binding prediction models that incorporate MS eluted peptide (EL) data, it could be useful to account for DO:DM and its correlation with abundant peptides that might otherwise be deemed weak binders. Such a model could, for example, be appropriately applied to naïve or memory B cells’ repertoires as opposed to activated B cells, which have high and low DO:DM, respectively. Relatedly, motif prediction, which is integral to binding prediction, also needs to consider the possible contribution of high DO:DM versus low DO:DM. We observed a swap of P1 dominant residue (F->Y) when DO:DM changes from infinity-high to medium-low-zero. A similar P1 residue swap was also found when comparing the EL-data-based motif versus BA-data-based motif. It is possible that DM editing selects for alternative high abundant registers between states.

Third, DO:DM tuning in APCs likely affects the length of presented peptides and generates longer peptide repertoires as the free DM levels increase. This hypothesis stems from the observation that longer peptides were presented in (DO:DM)^0-M^ as compared to DO^null^DM^null^ and (DO:DM)^H^ (Fig. 4E). More flanking residues can enhance MHC-II binding and increase DM-resistance (57, 58). For example, once a long peptide sits in MHC-II binding groove using a register biased towards the peptide’s C-terminus, additional N-terminal amino acids extending beyond the binding groove can create steric interference for DM access: DM associates with MHC-II from the N-terminal end of the loaded peptide (23, 59). Longer peptides could also provide more alternative binding registers. In addition, increased free DM likely allows peptide exchange in earlier, less acidic endosomal compartments where different/reduced processing capacity may result in longer peptides; these are compartments where DO, when present, effectively inhibits DM (17). The length changes mediated by DO:DM can also affect T cell recognition and peptide immunogenicity (60–62).

Last, DO:DM tuning in APCs may affect immunodominant T cell epitope selection from across a given protein’s sequence. One such source protein, TFRC, generated multiple DR4-binding core peptides belonging to 4 different major types (II-V). When free DM was expected to be low (DO^null^DM^null^ and (DO:DM)^H^) cells, we observed increased abundances of DM-sensitive cores and concomitant decreased abundances of DM-resistant cores all derived from the TFRC protein. The converse was true for DM-resistant cores with strong predicted binding affinity, when free DM was expected to be high in (DO:DM)^0-M^ cells. These changes are independent of TFRC abundance across these states and mostly consistent with the second pMHC-II presentation characteristic mentioned above. This observation leads us to speculate that upon antigen stimulation, DO-expressing APCs could experience similar DO:DM tuning state progressions until conditions are met for stable epitope presentation. As a result, a range of pMHC-II could be made available for T cell inspection over the course of APC maturation.

In conclusion, our study points to DO:DM tuning as an important factor that modulates pMHC-II presentation. Our observations were made possible by a model system designed to recapitulate the variable levels of DO and DM seen in different kinds of APCs with different differentiation states – states that regulate the balance between vigilance against pathogens versus tolerance of self. Accounting for multiple DO:DM states in the context of DO regulation will guide future investigations, such as comparing pMHC-II repertoires derived from multiple HLA-II alleles’ respect to varying DO:DM, and from primary cells that are sorted into different DO:DM states. In addition, further exploration of scenarios in which DO expression is limited (e.g., when macrophages or DO^null^ DCs present antigen to T cells) is warranted, because it is likely that we have also underestimated the graded effects of varied levels of DM can have on pMHC-II presentation. Experiments like these stand to promote improved models of MHC-II antigen presentation that can be generalized to a wider array of APCs with important relevance to human health.

## Abbreviations

DTT: Dithiothreitol
FDR: False discovery rate
HLA-II: Human leukocyte antigen class II
MHC: major histocompatibility complex
PBS: Phosphate buffered saline

## FUNDING

This work was supported by a Damon Runyon-Rachleff Innovation Award from the Damon Runyon Cancer Research Foundation (DRR-13-11; J.E.E), the W.M. Keck Foundation Medical Research Program (J.E.E.), and the Chan Zuckerberg Biohub (J.E.E.), a Knut and Alice Wallenberg Foundation Postdoctoral Fellowship (to N.O.), as well as the NIH (AI-095813; EDM and LNA) and the Lucile Packard Foundation for Children’s health (E.D.M).

## ACKNOWLEDGEMENTS

We wish to acknowledge all members of the Elias and Mellins Labs for helpful discussions during the preparation of this manuscript.

## DATA AVAILABILITY

In addition, all peptide data and mass spec raw data files have been deposited in the PRIDE Archive at www.ebi.ac.uk/pride/archive under accession number PXD024392.

## Notes

### Competing Interest Statement

The authors have declared no competing interest.

