## Supplementary Table and Figures for "Tuning DO:DM ratios modulates MHC class II immunopeptidomes"

#### SUPPLEMENTARY TABLES

| Sample | Unique peptide IDs | Unique peptide IDs (quant both bioreps.) | Cores | Cores for NetMHC |
|---|---|---|---|---|
| T2DR4 (biorep1)<br>T2DR4 (biorep2) | 2768<br>2873 | 1380 | 476 | 475 |
| T2DR4DM (biorep1)<br>T2DR4DM (biorep2) | 3119<br>3349 | 1943 | 527 | 525 |
| T2DR4DMDO(+) (biorep1)<br>T2DR4DMDO(+) (biorep2) | 3514<br>3594 | 2187 | 607 | 605 |
| T2DR4DMDO(++) (biorep1)<br>T2DR4DMDO(++) (biorep2) | 3337<br>3125 | 1839 | 538 | 534 |
| T2DR4DMDO(+++) (biorep1)<br>T2DR4DMDO(+++) (biorep2) | 3423<br>3529 | 1849 | 658 | 656 |
| T2DR4DMDO-Knockout (biorep1)<br>T2DR4DMDO-Knockout (biorep2) | 1502<br>2727 | 799 | 243 | 242 |
| Total # of unique peptides in dataset | 10587 | 4528 | 1206* | 1206* |

*Total core epitopes when all cell lines analyzed together

**Supplementary Table 1.**

General statistics of the MS-derived HLA-DR4 peptidome data, summarizing peptide and core identifications. In case of multiple peptide cores sharing an identical amino acid sequence but identified by MS to contain variable modifications (such as oxidized methionine, deamidation, cysteinylation or phosphorylation), only the sequence was included for NetMHCIIpan-4.0 analysis and enumerated once. A small number of cores containing unidentified amino acids denoted as “X” or “Z” were excluded from NetMHCIIpan-4.0 analysis.

**Supplementary Table 2. 1,206 peptide cores deduced from Plateau algorithm.** See file Supplemental_table_2.xlsx

**Supplementary Table 3. 729 peptide cores with significant variance across cell lines, and associated elution data, corresponding with Figure 4C.** See file Supplemental_table_3.xlsx

### SUPPLEMENTARY FIGURES

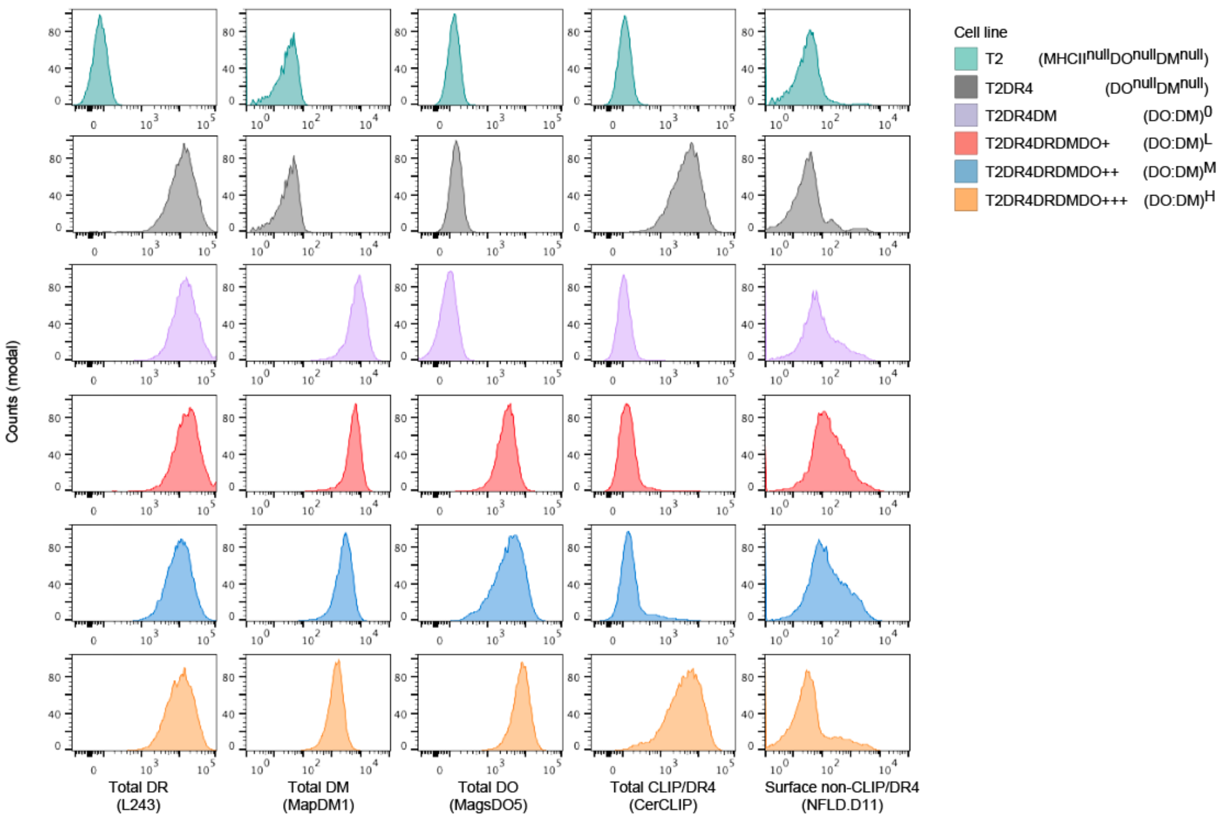


**Supplementary Figure 1. Representative flow cytometric histograms showing the expression of DR, DM, DO, CLIP/DR4 complexes and surface non-CLIP/DR4 complexes in different T2-derived cell lines.** Monoclonal antibodies (mAb) used for staining are L243, MapDM1, MagsDO5, CerCLIP, and NFLD.D11, as indicated. See Fig. 2B for quantification.

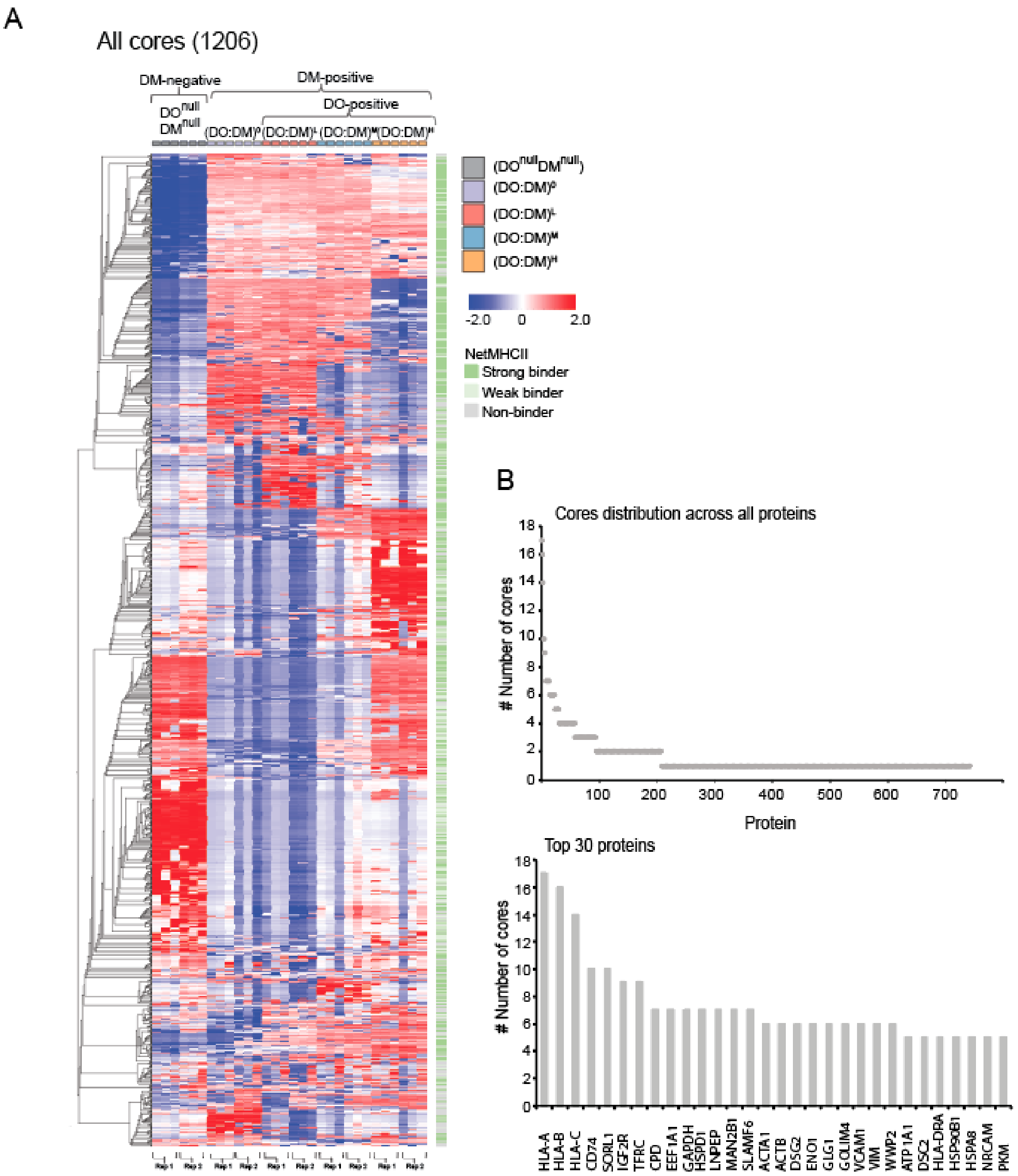


**Supplementary Figure 2. Identification of cores and their distribution across proteins.** (**A**) Heatmap illustrating all cores (z-score normalized). Analysis were limited to quantified peptides identified in both biological replicates. Cores predicted to be DR4-binders (top 10%rank, weak or strong) by NetMHCIIpan-4.0 are also depicted. **(B)** The distribution of binding cores across (top) all proteins and or (bottom) the 30 indicated proteins with highest number of total cores.

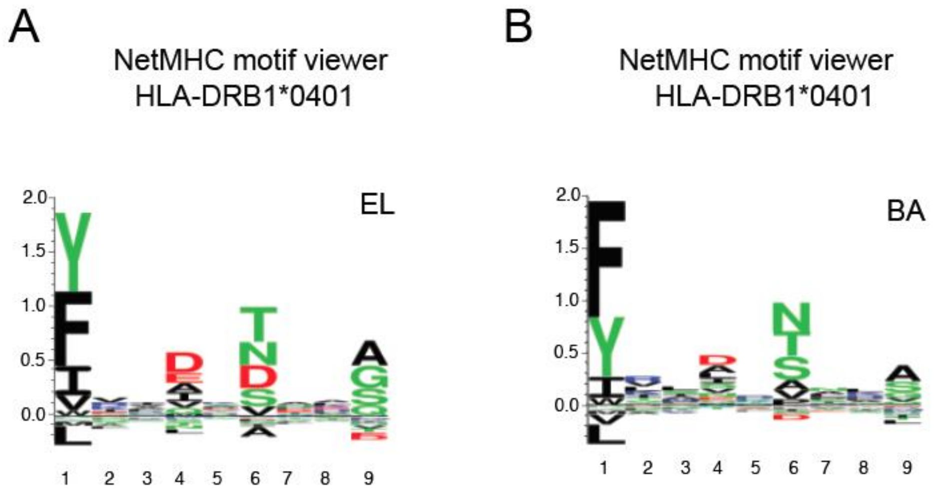


**Supplementary Figure 3. DR4 motifs downloaded from the NetMHCIIpan-4.0 motif viewer** (http://www.cbs.dtu.dk/services/NetMHCIIpan/logos.php). (**A**) Motif based on eluted Ligand mass spectrometry (EL) data **(B)** Motif based on Binding Affinity (BA).

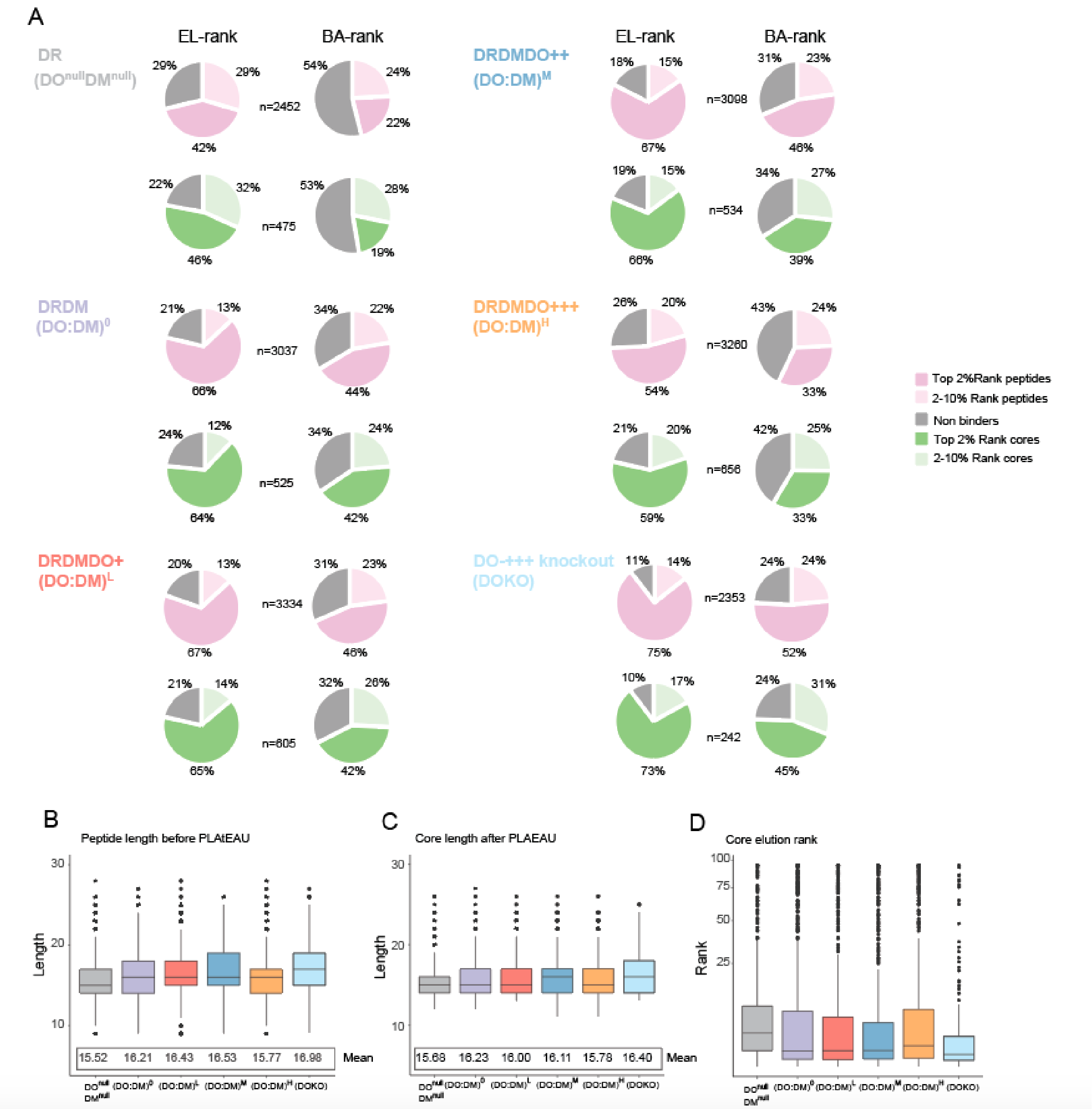


**Supplementary Figure 4. Evaluation of peptide and core epitope lengths, elution rank and binding rank affinities. (A)** Core epitope analysis through PLAtEAU was performed on an individual cell line level using all identified peptides per cell line. Average length values based on the number of identifications at the peptide and the consensus epitope level when using identified peptides for each cell line. Furthermore, binding affinity prediction results from NetMHC our demonstrated for both the elution rank score results and the binding assay scores. **(B)** Peptide length distribution of all identified peptides in each cell line. **(C)** Core epitope length distributions. **(D)** Elution rank score distribution for all core epitopes.

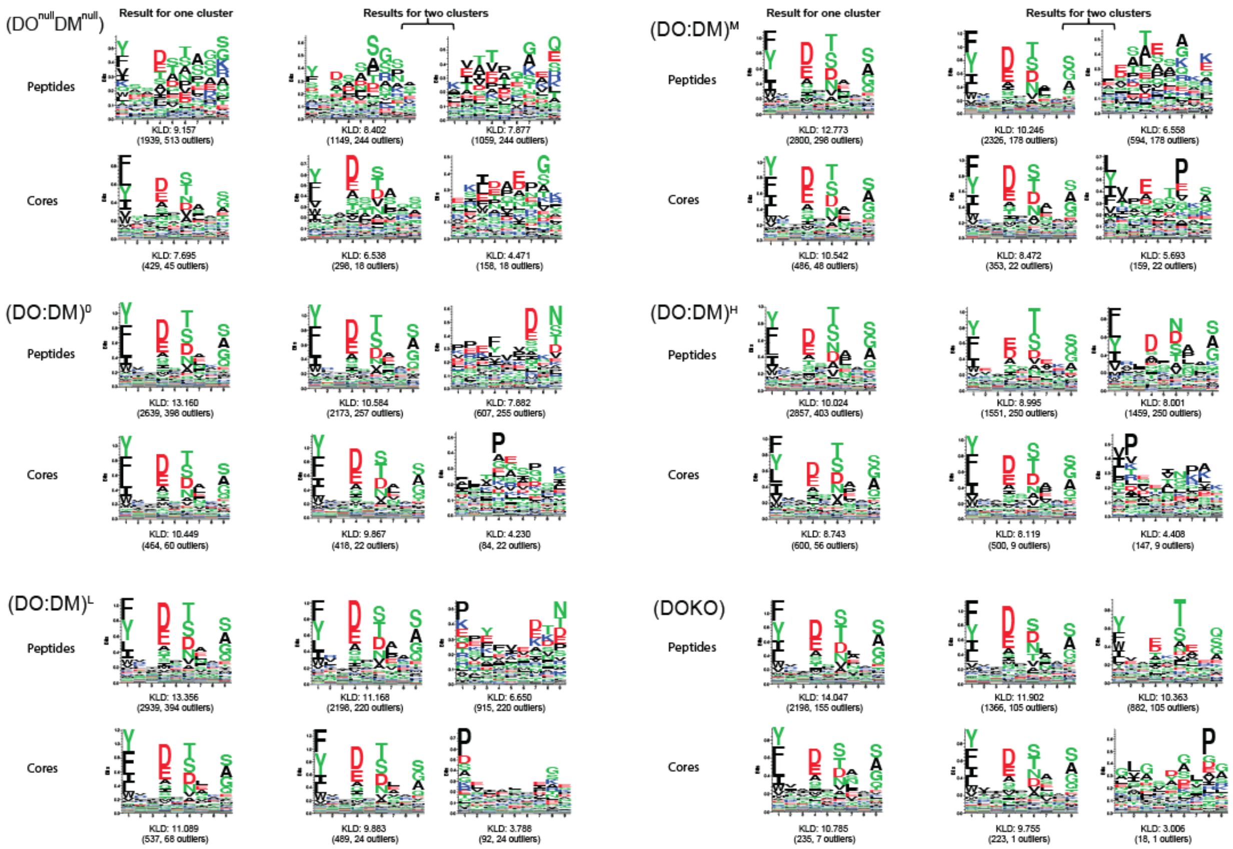


**Supplementary Figure 5.** Identification of DR4-like binding motifs using Gibbs cluster analysis for each individual cell line. Gibbs cluster analysis for all peptides or cores (through PLAtEAU) are demonstrated. These were illustrated using Seq2Logo. The top or top two reported clusters for each condition are presented. The Kullback-Leibler Distance (KLD) score is listed and the size of the cluster(s) and number of outliers in each cluster is listed in brackets.

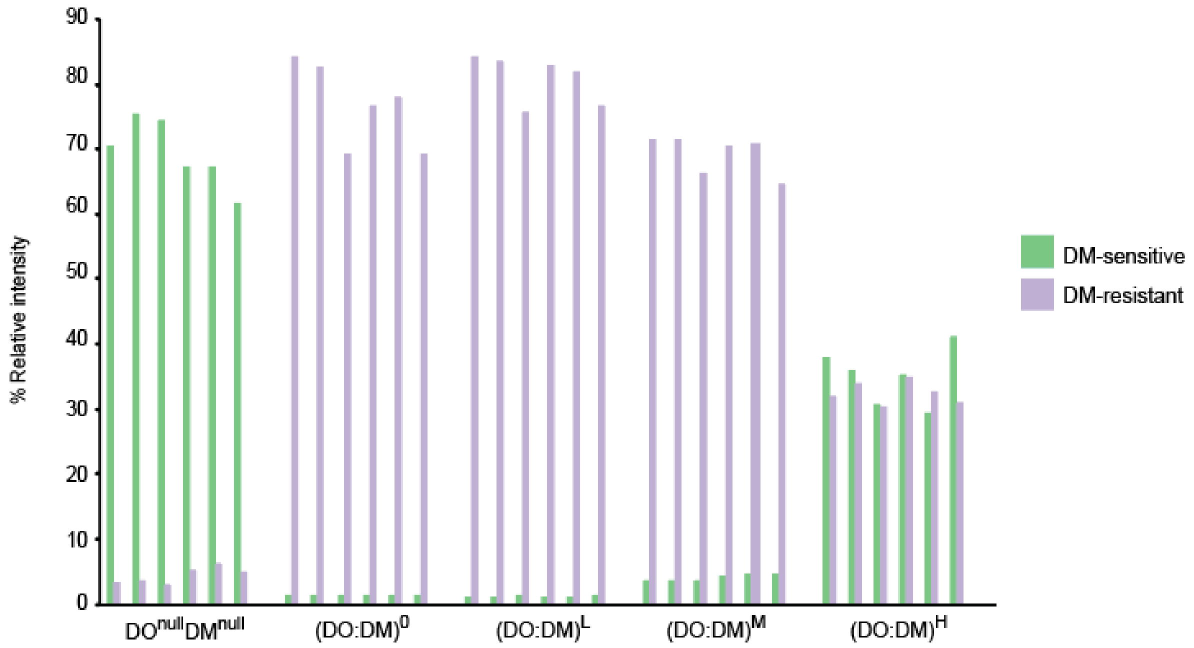


**Supplementary Figure**. **6**. % abundance subtotal of DM-sensitive (Series1=Type I-III) vs DM-resistant (Series2=Type IV-VI) cores (clustered in Fig. 4C) quantified in each technical replicate of different cell lines.

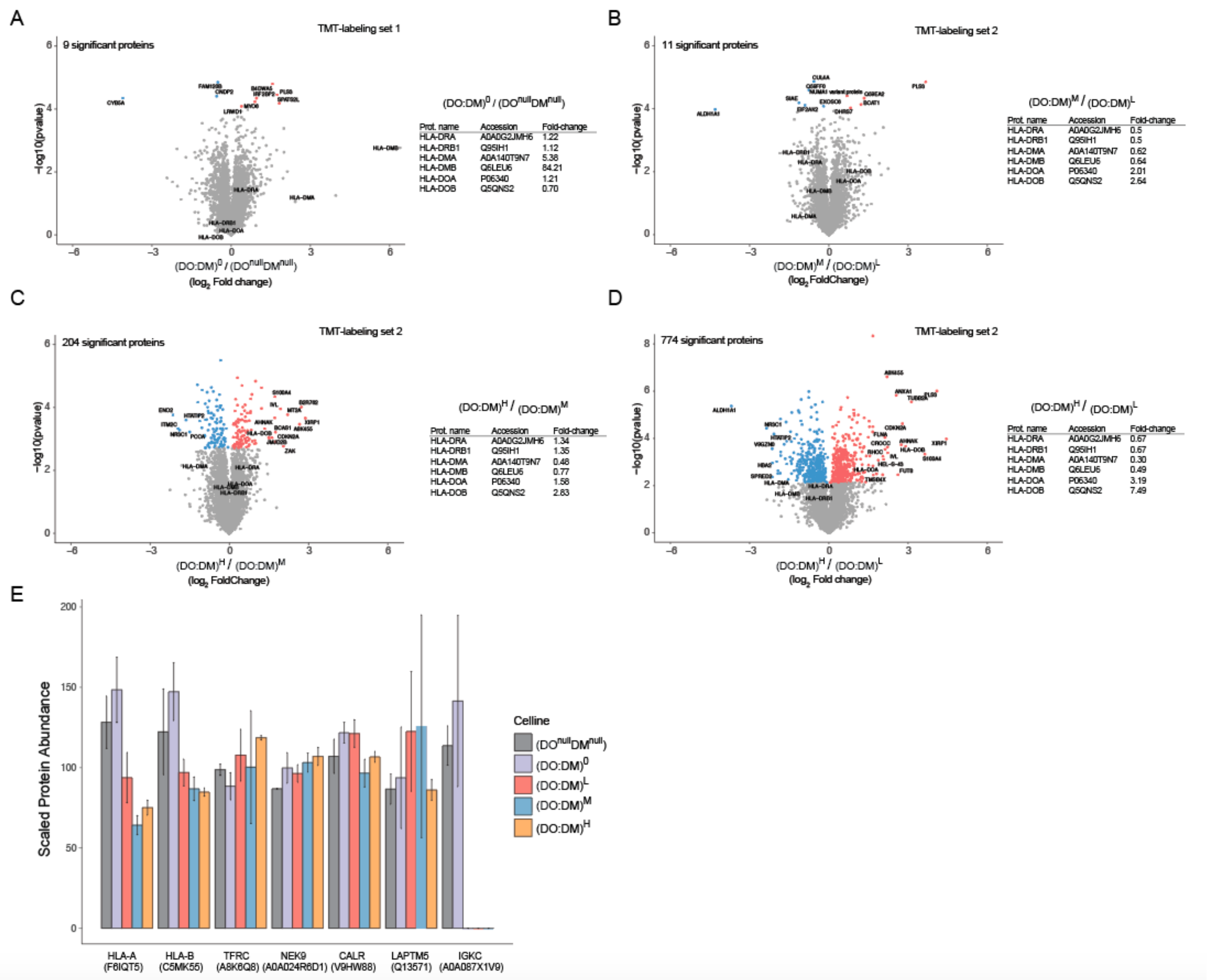


**Supplementary Figure 7.** Evaluation of differences in the proteomes between the DM-negative and DM-positive and the three DO-positive lines. **(A)** Minimal differential proteome expression differences were observed between the the $(DO:DM)^0$ and the DMnullDOnull line. Differentially expressed proteins (q-value of <0.05) that are upregulated indicated in red and downregulated indicated in blue. In addition to all the differentially expressed proteins a set of key proteins (including HLA-DRA, HLA-DRB1, HLA-DMA, HLA-DMB, HLA-DOA) are called out. **(B)** Proteome expression differences between the $(DO:DM)^M$ and the $(DO:DM)^L$ cell line. In addition to the differentially expressed proteins (q-value of <0.05) a subset of the key proteins (including HLA-DRA, HLA-DRB1, HLA-DMA, HLA-DMB, HLA-DOA) are called out. **(C)** Proteome expression differences between the $(DO:DM)^H$ and the $(DO:DM)^M$ lines. Differentially expressed proteins (q-value of <0.05) were all up-regulated indicated in red and downregulated are indicated in blue. In addition, to a subset of the most differentially expressed proteins, a selected set of proteins (including HLA-DRA, HLA-DRB1, HLA-DMA, HLA-DMB, HLA-DOA) are also called out. **(D)** Proteome expression differences between the $(DO:DM)^H$ and the $(DO:DM)^L$ lines. Differentially expressed proteins (q-value of $<0.05$) were all up-regulated indicated in red and downregulated are indicated in blue. In addition, to a subset of the most differentially expressed proteins, a selected set of proteins (including HLA-DRA, HLA-DRB1, HLA-DMA, HLA-DMB, HLA-DOA) are also called out. **(E)** Histograms represent scaled TMT-labeled protein signals from the corresponding proteins HLA-A, HLA-B, TFRC, NEK9, CALR, LAPTM5 and IGKC ± SD. The signal was scaled through the pooled “bridge” samples used in each TMT-label set (Fig. 1).

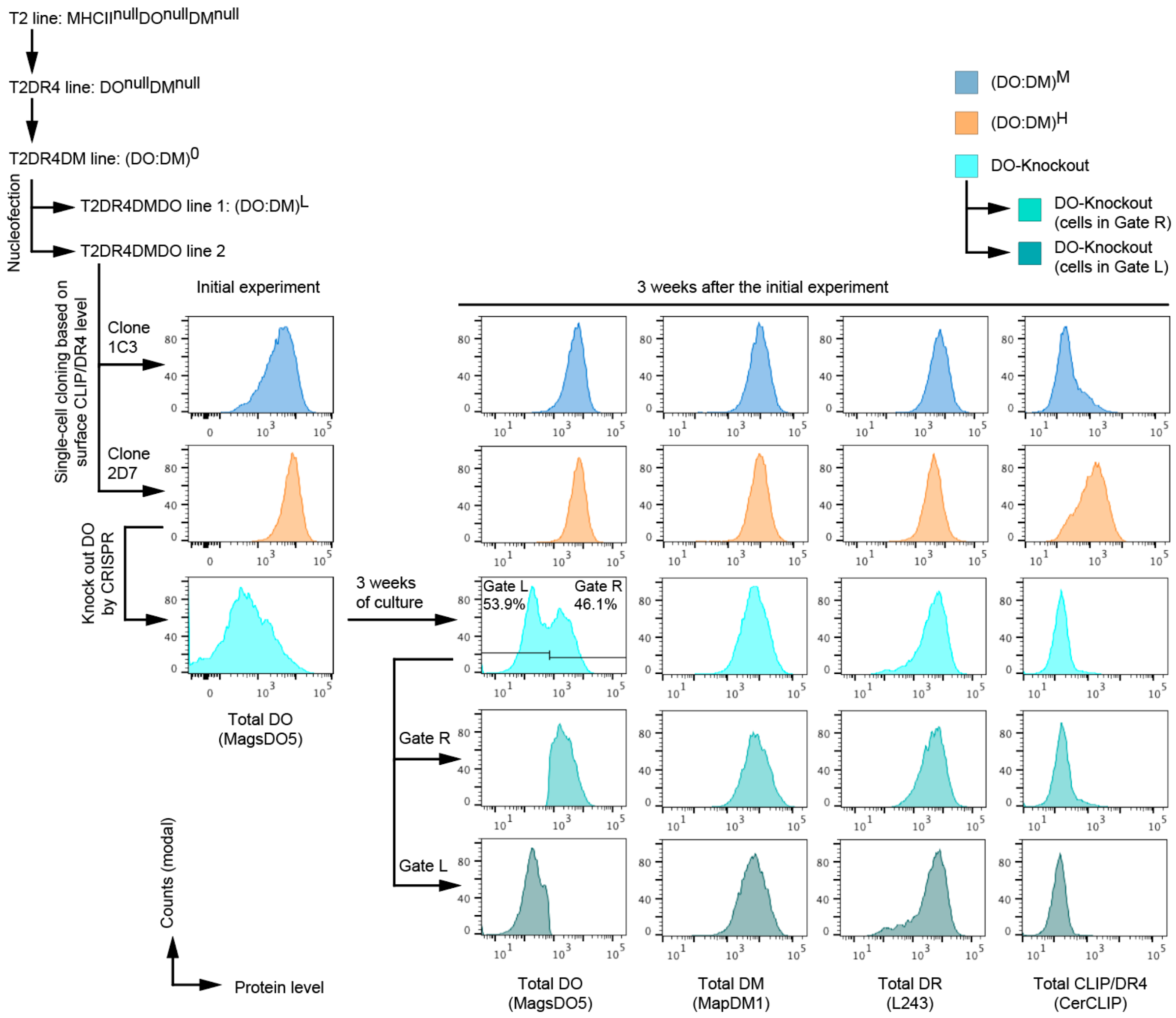


**Supplementary Figure 8. Flow data for DOKO to show varied DO expression levels in DOKO subpopulations.**

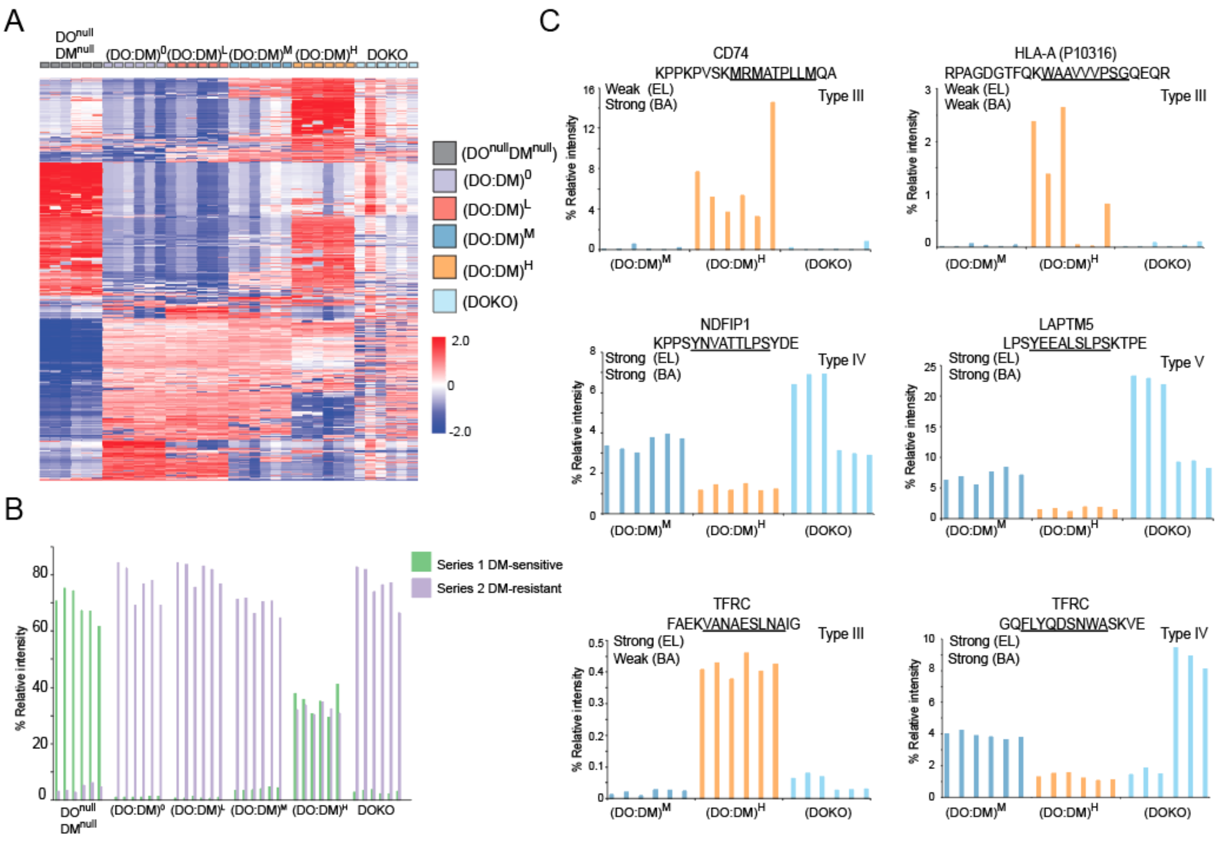


**Supplementary Figure 9. Reducing the levels of DO by CRISPR knockout results in a more $(DO:DM)^L$ like and typical DM-resistant phenotype**. **(A)** Heatmap (z-score normalized and ordered based on the hierarchy clustering established in 4C) with the addition of the DOKO line. **(B)** % abundance subtotal of DM-sensitive vs DM-resistant cores (clustered in Fig. 4C) quantified in each technical replicate of DOKO vs others. **(C)** Three examples of type III cores from CD74, HLA-A and TFRC. Two examples of Type IV cores for the NDFIP1 and TFRC protein. One example of a type V core from LAPTM5.

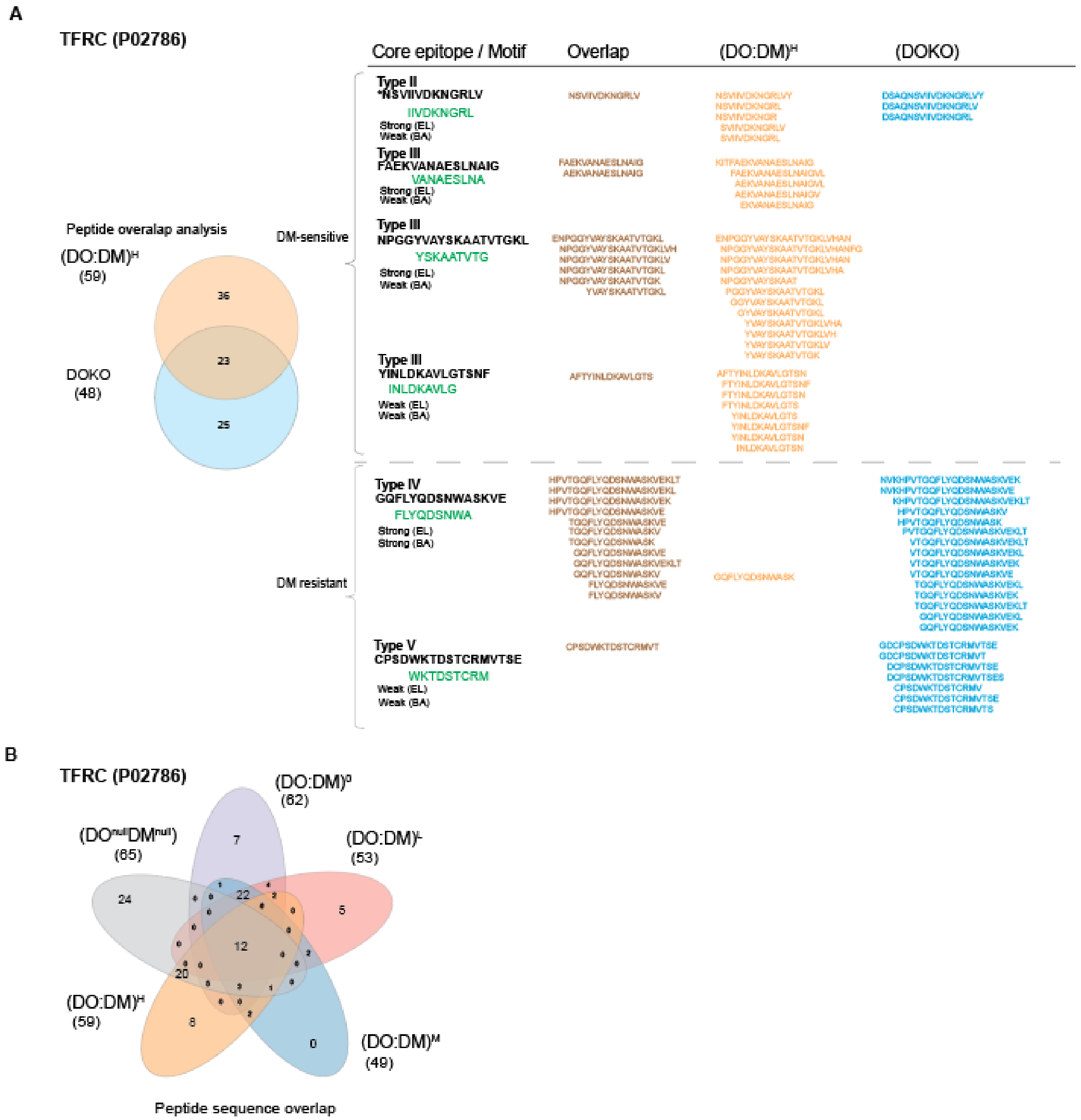


**Supplementary Figure 10. Different levels of DO reveals clear changes in TFRC cores and peptides. (A)** Direct comparison of peptides presented between the DOKO and the (DO:DM)$^H$ line. The peptides were matched up to the corresponding defined core types from figure 4C. **(B)**

Unique peptide sequence overlap analysis illustrated as a venn-diagram between the five lines excluding the DOKO.

#### SUPPLEMENTARY DATA

**Supplementary Data 1. Peptides and proteins identified and quantified from three TMT data sets spanning T2 and six T2-derived cell lines.** See file Supplemental_Data_1.xlsx

**Supplementary Data 2. Peptides identified and quantified from six T2-derived cell lines.** See file Supplemental_Data_2.xlsx
